# Meta-AlignNN: A meta-learning framework for stable brain-computer interface performance across subjects, time, and tasks

**DOI:** 10.1101/2025.04.20.649482

**Authors:** Yongjie Zou, Kai Liu, Congcong Zhang, Yu Ling, Fei Wang, Mingxin Li, Yulei Chen, Mengyu Li, Shouliang Guan, Zhenliang He, Chengyu T. Li

**Affiliations:** Lingang Laboratory, Xuhui District, Shanghai, 200031, China

**Keywords:** BCI stabilization, meta learning, alignment, neural dynamics, behavioral consistency, motor cortex

## Abstract

Practical application of brain-computer interfaces (BCIs) requires stable mapping between neuronal activity and behavior through various behavioral contexts and for different individuals. Due to neural activity instability, BCIs require frequent recalibration to maintain robust performance. Early approaches to addressing BCI stability issues mainly focused on tackling the challenge of neural activity changes over time. However, future BCI applications involve diversified scenarios and subjects, requiring solutions that address neural variability across time, subjects, and tasks. This study proposes a meta-learning-based algorithm to achieve BCI stability, named “Meta-AlignNN.” By capitalizing on the consistency of neural population dynamics, it provides a unified solution for maintaining BCI stability, robustness, and scalability across subjects, time, and tasks. Tested over two years on four tasks with three monkeys, as well as on public datasets, the approach has achieved significantly excellent performance in both offline decoding and real-time brain-control, outperforming existing methods. Our findings provides a foundation for meeting the clinical demands of longterm, efficient, and stable usability across patients and tasks, offering a compelling solution for practical BCI applications.

## Introduction

Brain-computer interfaces (BCIs) enable individuals with paralysis to control external devices by decoding neural signals. Noninvasive modalities including magnetoencephalography/electroencephalography (MEG/EEG) and functional magnetic resonance imaging (fMRI) offer low-risk neural signal acquisition, though constrained by inherent limitations in spatial (MEG/EEG) and temporal (fMRI) resolution (Buzsáki et al., 2012). In contrast, intracortical BCIs (iBCIs) demonstrate superior performance through high-resolution recordings (Willett et al., 2020) but face challenges from signal nonstationarities caused by electrode-related instabilities (impedance fluctuations, mechanical displacement, and failure) (Santhanam et al., 2007), tissue response dynamics (gliosis) (Chestek et al., 2011), and intrinsic neuronal variability (Churchland and Shenoy, 2007). These instabilities, occurring across minutes to weeks (Downey et al., 2018; Perge et al., 2013; Dickey et al., 2009), and variability across subjects/tasks, pose significant challenges for the practical application of BCIs.

Early methods addressing instability involved adaptive parameter updates (Orsborn et al., 2012; Dangi et al., 2013), tracking nonstationarities in neural recordings (Zhang and Chase, 2013; Bishop et al., 2014), dimensionality reduction (Yu et al., 2008; Shenoy et al., 2013; Nuyujukian et al., 2014; Sadtler et al., 2014), probability density function alignment (Dyer et al., 2017), and large-scale training strategies (Sussillo et al., 2016). As motor cortical activity during movement occupies a stable subspace (Churchland et al., 2012), leveraging low-dimensional neural dynamics further improved decoding stability (Kao et al., 2017; Pandarinath et al., 2018; Degenhart et al., 2020). Later work by (Gallego et al., 2020) systematically validated the long-term stability of movement-related manifolds, providing empirical justification for earlier algorithmic approaches.

Recent advances have integrated adversarial representation learning and generative models to address neural recording instabilities. By leveraging the Generative Adversarial Network (GAN) architecture (Goodfellow et al., 2014), ADAN was introduced to directly match empirical probability distributions of the latent variables across days (Farshchian et al., 2018). DANN combined feature extractors with gradient reversal layers to generate domain-invariant features, achieving cross-session generalization without labeled target data (Ganin et al., 2016). In terms of data augmentation, a generative model capable of synthesizing new data based on the learned mapping between hand kinematics and the corresponding neural spike trains was proposed by (Wen et al., 2023), enabling rapid adaptation with limited additional neural data. Unsupervised domain adaptation methods further minimized the need for daily labeled data (Jude et al., 2022; Degenhart et al., 2020).

Despite these advancements, critical challenges persist. Obviously, future BCI applications involve diversified scenarios and subjects, requiring solutions that address neural variability across time, subjects, and tasks. Recent studies (Gallego et al., 2018, 2020; Safaie et al., 2023) have emphasized the high preservation of neural patterns across subjects, time, and tasks, which inspired our proposed “Meta-AlignNN” and provided a theoretical foundation for its effectiveness. Meta-AlignNN is a novel meta-learning-based approach (Schmidhuber, 1987; Schaul and Schmidhuber, 2010; Tian et al., 2022) that combines a decoder trained on stable sessions with a meta aligner. The meta aligner is optimized using Model-Agnostic Meta-Learning (MAML) strategy (Finn et al., 2017) and is capable of aligning neural activity from different sessions into a consistent latent representation. This enables rapid adaptation to new sessions, subjects, and tasks with minimal data, akin to a “plug-and-play” approach, allowing new individuals or tasks to be integrated into existing BCI systems using only a subset of trials, thereby maintaining robust BCI performance amid neural drifts. By effectively capturing preserved latent dynamics underlying natural behavioral consistency, Meta-AlignNN provides a unified solution that maintains BCI stability, robustness, and scalability across subjects, time, and tasks using only a small number of stable recording channels, thereby significantly reducing the reliance on high-throughput recordings.

Validated on over 300 sessions (three monkeys, four tasks) and a public dataset (O’Doherty et al., 2017), Meta-AlignNN demonstrates significantly improved decoding performance compared to baseline methods, and achieves remarkable results in real-time brain-control across time, tasks, and subjects. Our findings provides a foundation for meeting the clinical demands of long-term, efficient, and stable usability across patients and tasks, offering a compelling solution for practical BCI applications.

## Results

### Instabilities of neural activity recordings across time, subjects, and tasks

Over approximately two years, three rhesus macaques performed four different tasks (see Methods; Figure 1a). Our self-developed flexible micro-electrode arrays were chronically implanted in primary motor cortex (M1) and dorsal premotor cortex (PMd) of monkeys, and neural activity along with the behaviors were recorded (see Methods). Neural recordings were converted into multiunit threshold crossing rates (see Methods). Neural signals recorded from implanted flexible micro-electrode arrays can maintain a high number of effective channels over an extended period, and neural recordings from some electrodes remain stable across multiple days (“Stable unit” in Figure 1b presents an example). However, at the same time, neural activity recorded from some other electrodes undergoes significant changes across days (Figures 1b and S1) and tasks (Figures 1d and 1e). More evidently, the neural signals recorded from corresponding channels in different monkeys also exhibit substantial differences (Figure 1c). Note that in Figure 1, the target directions for both js-random and wam-4 are involved. The target directions for js-random are explained in **Across-task paradigm design** method, while in wam-4, the target directions correspond to the positions of the moles (similar to up, down, left, and right in js-8). Similar with (Degenhart et al., 2020), instabilities observed in intracortical electrode recordings (lower part of Figures 1b and 1e) typically involve following situations across days and tasks: (1) baseline shift, a persistent alteration in an electrode’s baseline firing rate; (2) unit dropout, the irreversible loss of a neuronal unit’s detectable activity, caused by neuron death or physical displacement beyond the electrode’s detection range; (3) tuning change, the change of the functional relationship between neural firing patterns and behavioral variables, attributable to dynamic neuronal change and composition within the electrode’s recording vicinity; and (4) complex combination, any combination of the above three types. Across different subjects (lower part of Figure 1c), the instability is also similarly characterized by: (1) baseline difference, where there is a difference in the baseline firing rate of a corresponding channel pair across monkeys; (2) unit mismatch, where different sets of neurons are recorded on the corresponding channels across monkeys; (3) tuning difference, where there is a difference in the functional relationship between the neural firing patterns and the intended behavioral variables across monkeys; and (4) complex combination, any combination of the above three types. Distinct neural signatures were observed for each behavioral task (Figure 1d). As time, subjects, and tasks change, neural activity undergoes complex variations, resulting in low neural activity correlation across time, subjects, and tasks (Figure 1f). The correlation distributions for across-time, across-task, and across-subject scenarios each show significant differences compared to their respective within distributions. Additional details involved in Figure 1f are provided below: “within” refers to the firing rate correlation between the first and second halves of the same session on the same channel, while “across” refers to the firing rate correlation between corresponding channel pair across different sessions. Notably, for across-subject, “corresponding channel pair” refers to manually paired channels (from different monkeys but with similar implant locations), whereas for across-day or across-task, “corresponding channel pair” refers to the same physical channel.

**Figure 1.**
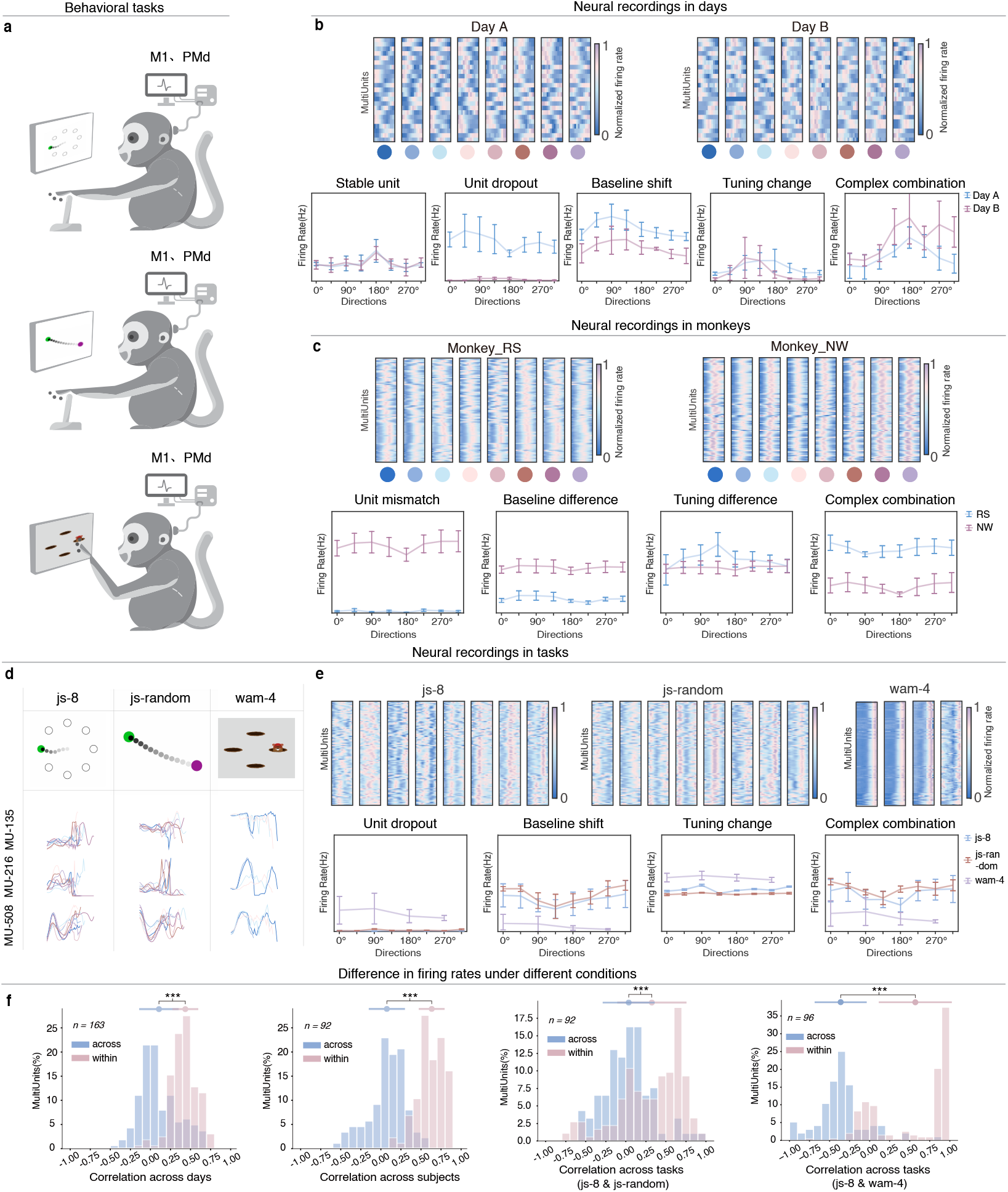
Variations in neural recording and activity patterns observed across days, subjects, and tasks. **a**, The monkey performed three tasks: 1) A center-out reaching task with eight targets (js-8, top panel); 2) A random target task (js-random, middle panel); and 3) A *whack-a-mole* task involving arm movements to touch four holes (wam-4, bottom panel). Flexible micro-electrode arrays were chronically implanted in the primary motor cortex (M1) and dorsal premotor cortex (PMd) to record neural activity during these behavioral tasks. **b**, Top: Example mean neural firing rates aligned to stimulus onset (23 recording channels over two days; vertically arranged by recording channel) and corresponding target directions of js-8 (bottom; eight colors are used to represent the eight adjacent targets in sequence). Columns represent trial-averaged responses for each of eight target directions. Obviously, neural activity underwent significant changes across days (divergent firing rates and tuning on each electrode), despite preserved behavioral consistency. Bottom: Representative directional tuning curve over two days in js-8. Each panel shows the single-electrode average firing rate profiles (error bars, mean ± s.d.) as functions of reach direction on two experimental days. Firing rates was quantified within a 1,200-ms epoch (-200 to +1,000 ms relative to stimulus onset). **c** and **e**, demonstrate similar trends across subjects and tasks, respectively. The target directions in js-random are explained in **Across-task paradigm design** method, while those in wam-4 correspond to the four cardinal positions of mole holes (analogous to the up, down, left, and right directions in js-8). **d**, These three recording channels illustrate the differences in firing rates between different units, as well as dynamic reorganization of neural patterns across behavioral tasks. Each curve represents the average firing rate of trials with the same target direction per recording session, with colors indicating different directions. **f**, From left to right, show the distribution of firing rate correlations per channel pair for across/within-day, across/within-subject, and across/within-task (js-8<->js-random, js-8<->wam-4), respectively. ***P<0.001 (two-sided Wilcoxon’s rank-sum test); n indicates the number of channel pairs; error bars represent mean ± s.d.

### Overview of Meta-AlignNN

Meta-AlignNN, somewhat similar to FA stabilization (Degenhart et al., 2020), ADAN (Farshchian et al., 2018) and SABLE (Jude et al., 2022), all of which leverage the premise that latent representations preserving movement intent remain stable despite nonstationarities in recorded neural activity. Recent studies have explored the stability and consistency of neural population dynamics from various perspectives, highlighting how these properties support the generation and learning of complex behaviors (Gallego et al., 2018, 2020; Safaie et al., 2023). These studies emphasize the high preservation of neural patterns across subjects, time, and tasks, forming the theoretical foundation for the effectiveness of Meta-AlignNN.

Meta-AlignNN is built on the principles of metalearning to identify and preserve invariant neural representations that exhibit stability across subjects, sessions, and behavioral tasks. We developed a meta aligner capable of aligning diverse neural activity to stable and consistent neural latent dynamics across subjects, time, and tasks, while being robust to changes or drift in neural activity recorded by cortical electrodes. (Figure 2a). Meta-AlignNN adopts a modular architecture consisting of a meta-learning-based alignment module and a fixed decoder. Taking js-random as an example (see Figure S2a), this two-stage framework operates through: (1) Alignment stage: Neural activity is mapped by the meta-aligner to a stable latent representation; (2) Decoding stage: A pretrained decoder is used to predict continuous estimates for BCI control. The parameters of the meta aligner adapt automatically across time, subjects, and tasks to mitigate recording instabilities, while the parameters of the decoder remain fixed.

**Figure 2.**
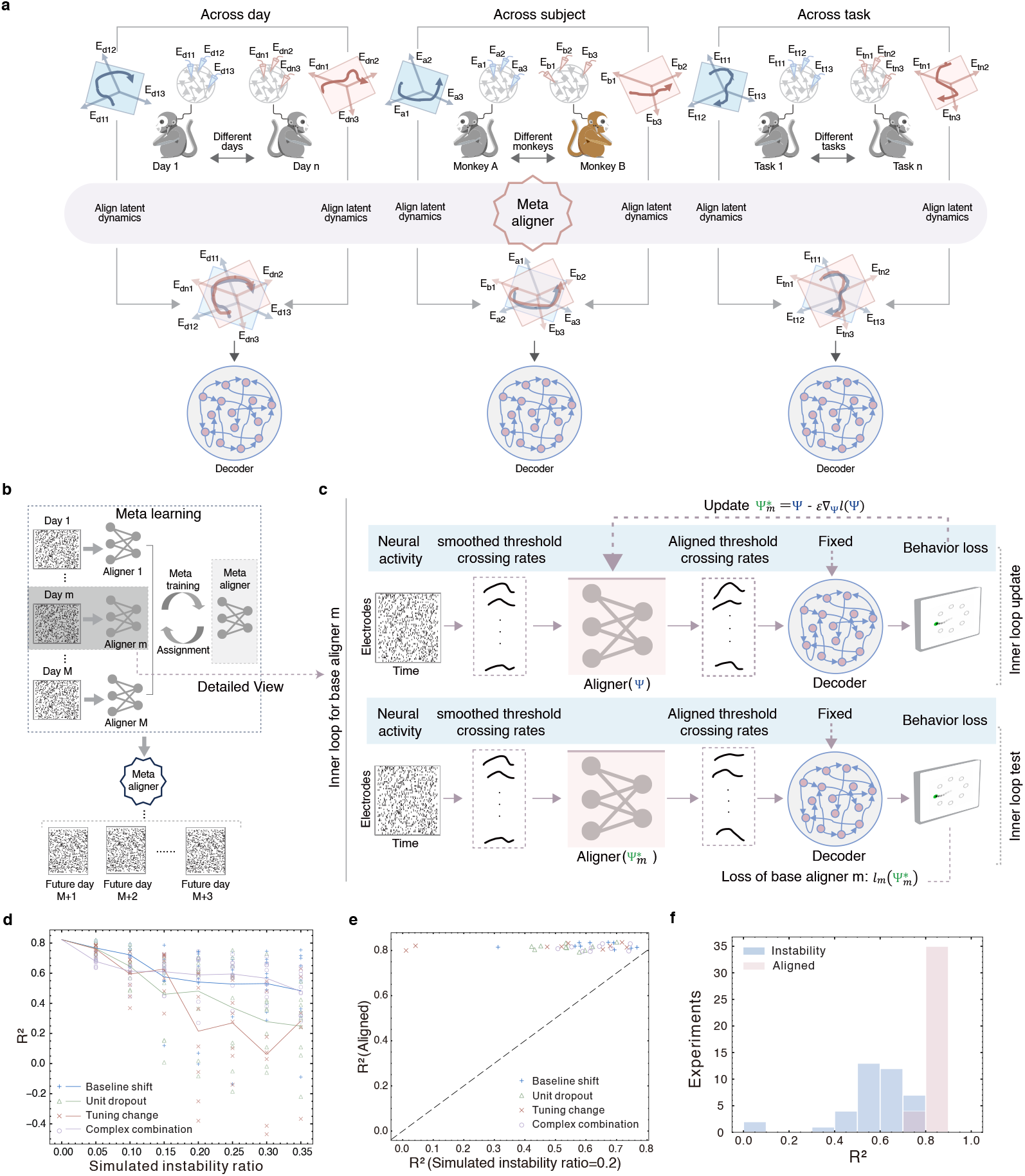
Overview of Meta-AlignNN and its effectiveness in handling simulated instability. **a**, Across-day: subjects perform the same task over days; Across-subject: different subjects perform the same task; Across-task: subjects perform various tasks. The meta aligner can effectively align latent dynamics in all three scenarios, enabling decoding stability across days, subjects, and tasks. **b** and **c**, Taking the across-day scenario as an example, here we show the model architecture of Meta-AlignNN and corresponding data flow processing. The overall structure and process remain consistent for across-subject and across-task scenarios. The differences lie in the definition and partitioning of training data, which are detailed in supplementary figure S2. **b**, illustrating the outer loop of training the meta aligner (meta-training process). Each day, a base aligner must be trained to align the neural data for that specific day. The parameter updates of the meta aligner depend on all the base aligners. Once the meta aligner is trained, it will be applied to future testing sessions. **c**, The loss computation of task *m* is illustrated for example. The upper half shows the process that update the meta aligner to adjust to task *m* through back propagation on behavior loss computed on its training data, while meta loss of task *m* is computed with input of test data in *m* that displayed in underneath half. The optimization objective of meta aligner is the sum of the meta losses across all *M* tasks. **d**, The relation between decoding performance degradation and perturbed ratio of different instability type. X-axis denotes the ratio of channel that applied with simulated instability. Each line color represent specific instability type and each dot represents individual experiments. **e**, For each experiment, decoding performance was quantified via coefficient of determination (R-squared) comparisons between aligned and instability evaluation. Different symbols represent different types of instabilities. Data points positioned above the dashed unity reference line indicate experiments where Meta-AlignNN enhanced decoding performance. Conversely, points below this line reflect conditions where Meta-AlignNN even had a counterproductive effect. **f**, Histograms of the R-squared for instability (blue) and alignment (claret) evaluation experiments.

#### The decoder

Strictly speaking, the term “decoder” here does not refer to a traditional decoder, as it consists of a Gated Recurrent Unit (GRU) encoder and a regressor. For simplicity, we refer to these two components collectively as the “decoder.” We trained the decoder using data from multiple high-quality recording days that were nearly consecutive, with one session per day. All sessions and electrodes used for training are carefully identified (see Methods). The type of decoder is not restricted theoretically and can be selected arbitrarily, e.g., Kalman filter (Wu et al., 2006; Wu and Hatsopoulos, 2008; Gilja et al., 2012), Naive Bayes (Barbieri et al., 2005; Kloosterman et al., 2014), Wiener filter (Carmena et al., 2003), Wiener cascade (Pohlmeyer et al., 2007), Support vector regression (Smola and Schölkopf, 2004; Chang and Lin, 2011), XGBoost (Natekin and Knoll, 2013), Feedforward neural network (Orbach, 1962; Goodfellow et al., 2016), Simple RNN (Rumelhart et al., 1986; Goodfellow et al., 2016), Gated recurrent unit (Cho et al., 2014; Goodfellow et al., 2016). In this study, we chose GRU as the base architecture for the decoder primarily because it is a nonlinear model capable of effectively capturing the stable latent representations embedded in neural activity. Notably, the initialization state of the decoder has a substantial impact on the performance of the meta aligner. The entire process can be formally described as follows: *z* = *g*(*x*), encoding neural activity *x* into latent variables *z*; *y* = *f* (*z*), decoding latent variables *z* into behavioral outputs *y*.

#### The meta aligner

Our approach is to learn an optimal mapping function *x*^∗^ = *ψ*(*x*), which could dynamically map non-stationary neural signals to stable latent dynamics, thereby facilitating behavioral decoding: *x*^∗^ = *ψ*(*x*), *z* = *g*(*x*^∗^), *y* = *f* (*z*). However, the alignment mapping *ψ* exhibits significant session dependency. Across different times, subjects, or tasks, variations in the recorded neural population and substantial changes in neural activity typically result in *ψ* taking on domain-specific transformation forms. To address this, we introduce the concept of meta-learning to discover the function *ψ*, and propose the notion of a “meta aligner” aimed at enhancing the generalization capability of the process of learning *ψ* itself.

During the training of the meta-aligner (see Methods), 80% of the sessions recorded on subsequent days (one session per day) were assigned as training tasks, and the remaining 20% as test tasks. It is important to note that the term “task” here does not refer to the four experimental paradigms, but instead treats each session as a task, with training and test data further partitioned within each task (Figure S2c). We trained the meta-aligner using MAML policy (Finn et al., 2017), during which the decoder remained fixed and only the parameters of the meta aligner were updated (e.g., in the cross-time scenario, Figure 2b,c, see Methods). The overall training procedure for the cross-subject and cross-task settings followed the same structure as the cross-time case, with the primary difference lying in the definition of tasks and the data partitioning scheme (Figure S2b,c).

The various types of recording instabilities naturally involved in the training tasks, such as baseline shift, unit dropout, tuning changes, complex combinations and other unobserved variability (Figure S2d), compelling the meta aligner, under the guidance of the meta-training mechanism, to learn how to generalize the alignment function *ψ* across diverse and complex scenarios. In fact, the substantial differences in time, subject, and task between the sessions used for early decoder training and those used for later meta aligner training (Figure 1 and S1) ensure the “difficulty” of the training tasks, thereby promoting the generalization ability of the meta aligner. We find that, provided sufficient training sessions (tasks), meta aligner can make fast adaptation to unseen sessions with merely minutes of neural data, facilitating rapid calibration to preserve control stability during severe signal quality deterioration.

Figure S2e provides an intuitive illustration: through meta-training, *ψ* is updated into a well-initialized parameter representation that can quickly adapt to the corresponding optimal parameters *ψ*^∗^(*instabilities*) when encountering a specific type of signal instability. From a data perspective, Figure S2f provides an intuitive illustration of the effect of *ψ*^∗^(*instabilities*).

#### The effect of transformer and meta training

Since transformer-based architecture of meta aligner and meta-training strategy are two important components for Meta-AlignNN, we conducted ablation studies to investigate their individual contributions to model performance. We compared variants of our model under the following two conditions: 1. Meta-AlignNN-Forward: The Transformer-based architecture (Vaswani et al., 2017; Devlin et al., 2018) is removed, and the meta aligner is replaced with a standard feedforward neural network. 2. AlignNN-No-Meta: Meta-learning is disabled, and instead, the session data used by the meta aligner are directly used for conventional training. As shown in Figure S3, evaluated R-squared as a function of the number of training epochs during inference stage, the results reveal the following: 1. AlignNN-No-Meta achieved worse performance compared with Meta-AlignNN under the same limited number of training epochs, which indicates that the necessity of meta training for aligner to take a quick adaptation to new neural recordings. 2. Meta-AlignNN clearly outperforms Meta-AlignNN-Forward under the same number of training epochs, implying that transformer is good at learning and capturing more powerful and generalized representation, even in the BCI domain.

These ablation studies strongly support the importance of the two core design choices in Meta-AlignNN: the Transformer architecture enhances the model’s representational capacity, while the meta-training strategy ensures its adaptive generalization capability.

### Meta-AlignNN demonstrates effectiveness in handling simulated instability

As mentioned above, instabilities observed in intracortical electrode recordings (Figure 1) typically can be summarised as baseline shifts/difference, unit dropout/mismatch, tuning changes/difference and complex combinations (Figure S2d). We specifically focus on evaluating Meta-AlignNN’s ability to handle these types of instabilities. First, We aimed to investigate the relation between decoding performance degradation and perturbed ratio of different instability type. A decoder was trained on the training set from a single session and then used to evaluate a test set from the same session with simulated instabilities introduced (as shown in Figure S2d). As shown in Figure 2d, decoding performance decreased nearly linearly with the rising of perturbed ratio. Among the perturbation types, baseline shift had the least impact on decoding, while tuning changes were the most detrimental. Next, we conducted 30 independent contrast experiments to evaluate the decoding performance for each type of simulated instability under two conditions: with and without alignment (i.e., using or not using Meta-AlignNN). The goal was to quantify Meta-AlignNN’s effectiveness in counteracting each of the four instability types. As shown in Figure 2e, Meta-AlignNN significantly improved R-squared under all instability types, with gains reaching up to approximately 0.8. Notably, all experimental dots are above diagonal line, indicating that Meta-AlignNN provided consistent positive improvements in every case. Figure 2f presents a histogram summarizing the results across all experiments (covering all four instability types), further confirming Meta-AlignNN’s robustness and generalizability in the face of diverse simulated neural perturbations.

### Meta-AlignNN is effective in addressing natural instability

We systematically evaluated the behavioral decoding performance of Meta-AlignNN and compared it with several state-of-the-art approaches on held-out neural recording sessions. The comparison methods (see Methods) include: FA stabilization (Degenhart et al., 2020), ADAN (Farshchian et al., 2018), SABLE (Jude et al., 2022), and DANN (Ganin et al., 2016). Additionally, we evaluated a baseline GRU, trained to decode movement kinematics directly from neural activity without meta aligner. To assess the effectiveness of Meta-AlignNN, we tested it on two types of reaching tasks: the js-8 task and a self-paced reaching task without gaps or pre-movement delay intervals (from a public dataset (O’Doherty et al., 2017)). For both tasks, we adopted the same evaluation strategy: all models were trained using historical training data and evaluated on subsequent held-out test sessions for behavioral decoding (i.e., 2D cursor position prediction), using only a small amount of neural data during testing for alignment. Implementation details for comparison models and Meta-AlignNN in this contrast experiments are provided (see Methods). The results, summarised in Table 1, show that Meta-AlignNN outperforms all comparison models in every case.

**Table 1.**
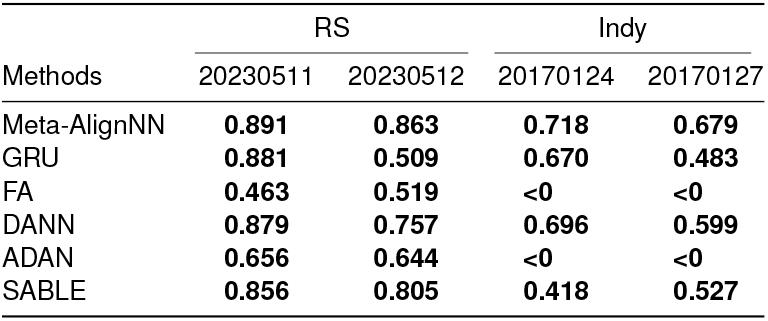
Methods and metrics. Each method was evaluated on two tasks that performed by two monkeys respectively. For each task, we test neural recordings of 2 sessions separately. The predictive accuracy was quantified as the mean R-squared between decoded and actual two-dimensional coordinates (x, y).

Meta-AlignNN demonstrated its performance in real-world across-time, across-subject, and across-task scenarios. Specifically, across-subject refers to using the decoder from one subject (Monkey_RS or Monkey_NW) to decode the behavior of the other subject in the js-8 task; across-task refers to using the decoder from js-random, js-4, and wam-4 tasks to respectively decode js-8. Compared to the unaligned condition, Meta-AlignNN significantly improved behavioral decoding performance for js-8 across time, subjects, and tasks (Figure 3). Figures 3a,d,g illustrate the input threshold crossing rates, the actual and predicted cursor positions, over eight randomly sampled test trials from session data across time, subjects, and tasks. Figures 3b,e,h summarize results from two test sessions: the first row the ground truth movement trajectories, and the second/third row represents the decoded reconstruction of movement trajectories without/with alignment respectively. Only a small number of trials were used to quickly adapt the meta-aligner for decoding the remaining unseen test data. In both sessions, the severe degeneration of decoder was occurring and the alignment with meta aligner compensate for the weakness that shows good overall reconstruction of movement trajectories. Figures 3c,f,i illustrate the quantify progressive enhancements in the meta aligner’s adaptation performance relative to increasing training data availability, assessed across time, subjects and tasks respectively. We add 10 seconds of training data every time to construct different number of training data used for meta aligner, and calculating the mean percentage increase in performance specifically for the last 25 percent of each test sessions. With very limited training data, prediction performance improved significantly, and the improvement trends were highly consistent and convergent across time, subjects, and tasks, which is ideal for practical applications.

**Figure 3.**
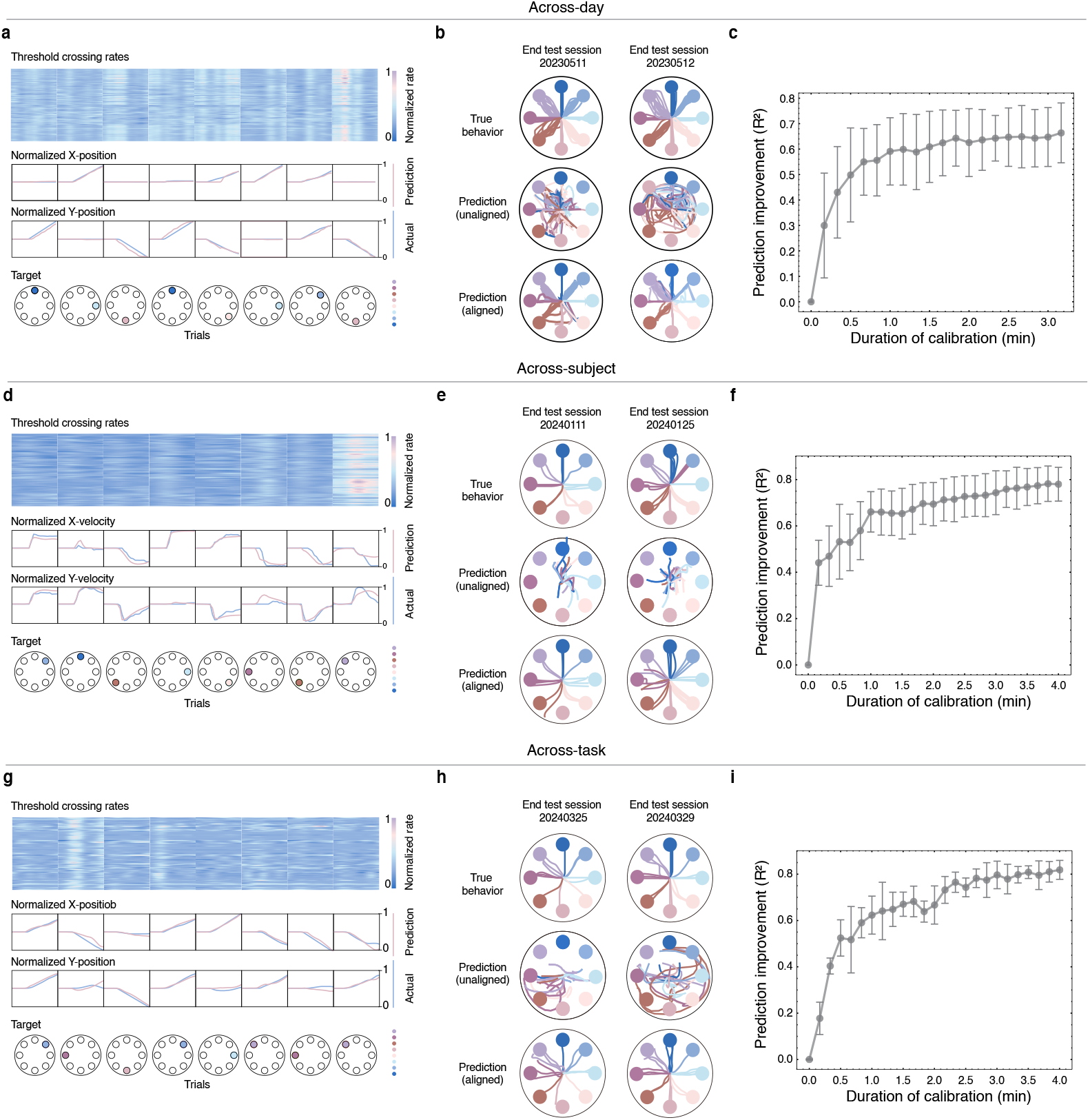
Meta-AlignNN demonstrates effectiveness in addressing natural instability across time, subjects, and tasks. **a**, Threshold crossing rates recorded from M1 and PMd over eight example trials while the monkey performed js-8 task. Trajectory prediction quality was evaluated through kinematic decomposition of two-dimensional cursor paths, with quantitative comparisons between decoded and actual positions across horizontal (X-axis) and vertical (Y-axis) components for eight representative movement trials. **b**, Comparisons between 2D monkey cursor position of test trials for using alignment or not in 2 sessions respectively, where a few data is used for alignment and subsequent end part is tested for decoding movement trajectories. **c**, The mean improvement in behavior prediction R-squared achieved through the use of the meta aligner for alignment, plotted against the quantity of training data required for the meta aligner’s adaptation at the start of each day, averaged across all test sessions (error bars, mean ± s.d.). Similar to **a, b**, and **c, d, e**, and **f** describe across-subject scenarios, while **g, h**, and **i** describe across-task scenarios.

### Meta-AlignNN enables neural activity alignment for consistent latent representations and improved behavioral decoding

Supplementary analyses reveal that Meta-AlignNN not only supports “good” CEBRA-Behavior embedding (Figure S4) but also enforces latent space consistency (Figure S5). In terms of neural latent trajectories and representations, we conducted further analyses using Meta-AlignNN for alignment across days, tasks, and subjects. As illustrated in Figures 4a1,a2,a3, two-dimensional t-SNE visualizations of trial-averaged latent trajectories are presented (see Methods), with each trajectory computed across all trials per target direction, in which all trials are aligned to movement onset. Figures 4a1,a2,a3 mean unaligned test session, past train session and aligned test session respectively. The differences between latent trajectories of past sessions and unaligned test sessions reflect neuronal turnover and recording instability. Comparison between trajectories in Figure 4a1 and Figure 4a2 identifies a suite of spatial transformations—shear distortions, rotation, and scaling. However, the similarity between trajectories in Figure 4a3 and Figure 4a2 proves that powerful mechanism of Meta-AlignNN bring us a good alignment. Figure 4b and Figure 4c quantify the improvement in neural latent representations brought by alignment through Meta-AlignNN. Figure 4b shows a comparison of the canonical correlations (CCs) of neural latent variables (see Methods). Across multiple test sessions, the pairwise CCs of unaligned neural latent variables are significantly smaller than those of aligned neural latent variables. Figure 4c reveals a comparison of the direction disorderliness of neural latent trajectories (see Methods). Across multiple test sessions, the direction disorderliness of unaligned neural latent trajectories is significantly larger than that of aligned neural latent trajectories, and the aligned condition is close to the within-condition. Similar to Figure 4a, the only difference in Figure 4d is that it shows the 2D t-SNE visualization across subjects and the only difference in Figures 4g,j,m is that they display the 2D t-SNE visualizations under three different across-task scenarios. “Monkey_RS -> Monkey_NW” means that Monkey_RS uses the js-8 decoding model from Monkey_NW to decode its own js-8. Other across-subject identifiers follow similarly. “Js-random -> js-8” means that the js-8 decoding model is used to decode js-random. Other across-task identifiers follow similarly. Similar to the across-time scenario, both CCs (Figures 4e,h,k,n) and direction disorderliness (Figures 4f,i,l,o) also show significant differences in the across-task and across-subject scenarios. However, all these trajectories are drawn by dimension reduced representations, which means that dimension reduction may bring one additional impact factor. As a result, to exclude this influence, 32D latent variables (with/without alignment) are shown in Figure S6a. In detail, Figure S6b illustrates correlations for latent variables of eight target directions respectively. Clearly, the correlation with alignment is significantly higher than the correlation without alignment. Finally, we want to find out correlation between latent variables of different targets with alignment. Correlation matrix is computed and visualized in Figure S6c. Interestingly, for eight targets, more spatially close to each other, corresponding latent variables after alignment are more similar. For example, as eight targets are distributed in ring structure, target 1 is closest to target 2 and 8, then target 3 and 7, next target 4 and 6, last target 5. The correlation between the latent variables of target 1 and the other targets follows the same order as the spatial sequence, which can be easily observed and concluded from Figure S6c.

**Figure 4.**
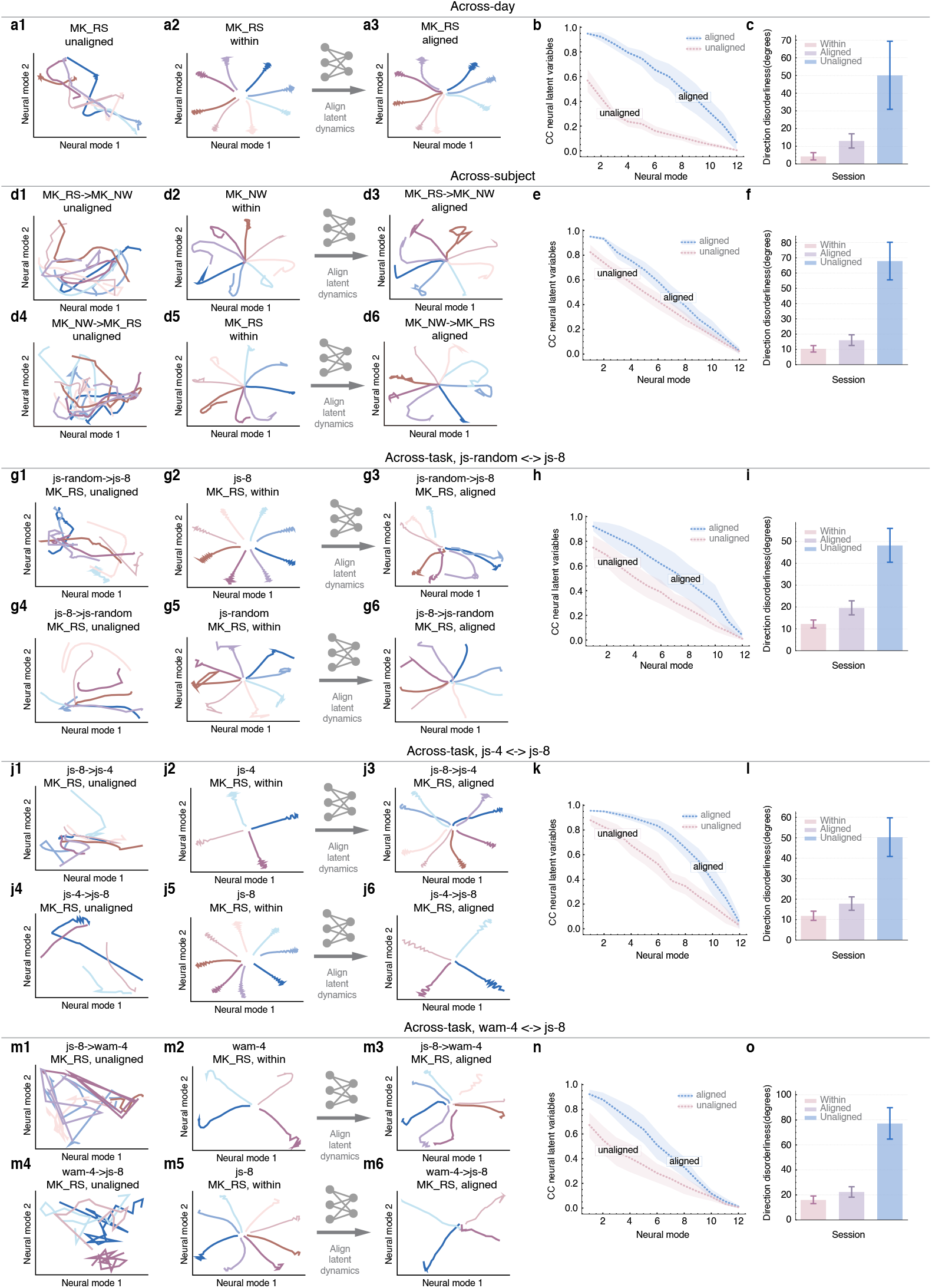
Comparison of neural latent trajectories with and without alignment. **a1, a2** and **a3** display two-dimensional t-SNE visualization of mean neural latent trajectories as the monkey performed js-8 for test session before alignment, within and after alignment respectively. **b**, Canonical correlations (CCs) analysis for neural latent variables. Line and shaded area, mean ± s.d. A marked decrease in pairwise CCs magnitudes was observed between the unaligned relative to aligned neural data. **c**, Comparison of direction disorderliness for neural latent trajectories. The direction disorderliness were significantly larger for the unaligned than for the aligned neural latent trajectories. Error bars, mean ± s.e. The aligned condition is already close to the within condition. Similarly, **d** to **f** show the situation across subjects (Monkey_RS<->Monkey_NW); **g** to **i** show the situation across tasks (js-random<->js-8); **j** to **l** show the situation across tasks (js-8<->js-4); **m** to **o** show the situation across tasks (js-8<->wam-4).

### Small number of stable multiunits is enough for Meta-AlignNN across days

We investigated whether Meta-AlignNN can maintain strong performance over long timescales after the decoder has been trained. Figure 5a summarizes the performance comparison of the fixed decoder before and after alignment by the meta aligner, quantified by the R-squared value. Each indigo symbol indicates the performance achieved with alignment in each individual experiment on each different day. Similarly, claret dot denotes using decoder directly without alignment. The claret line quantifies progressive performance degradation of non-adaptive decoders attributable to non-stationary neural signal drift across days, whereas the indigo line demonstrates performance improvement achieved via meta aligner. Remarkably, the improved performance was still largely maintained even after over 300 days.

**Figure 5.**
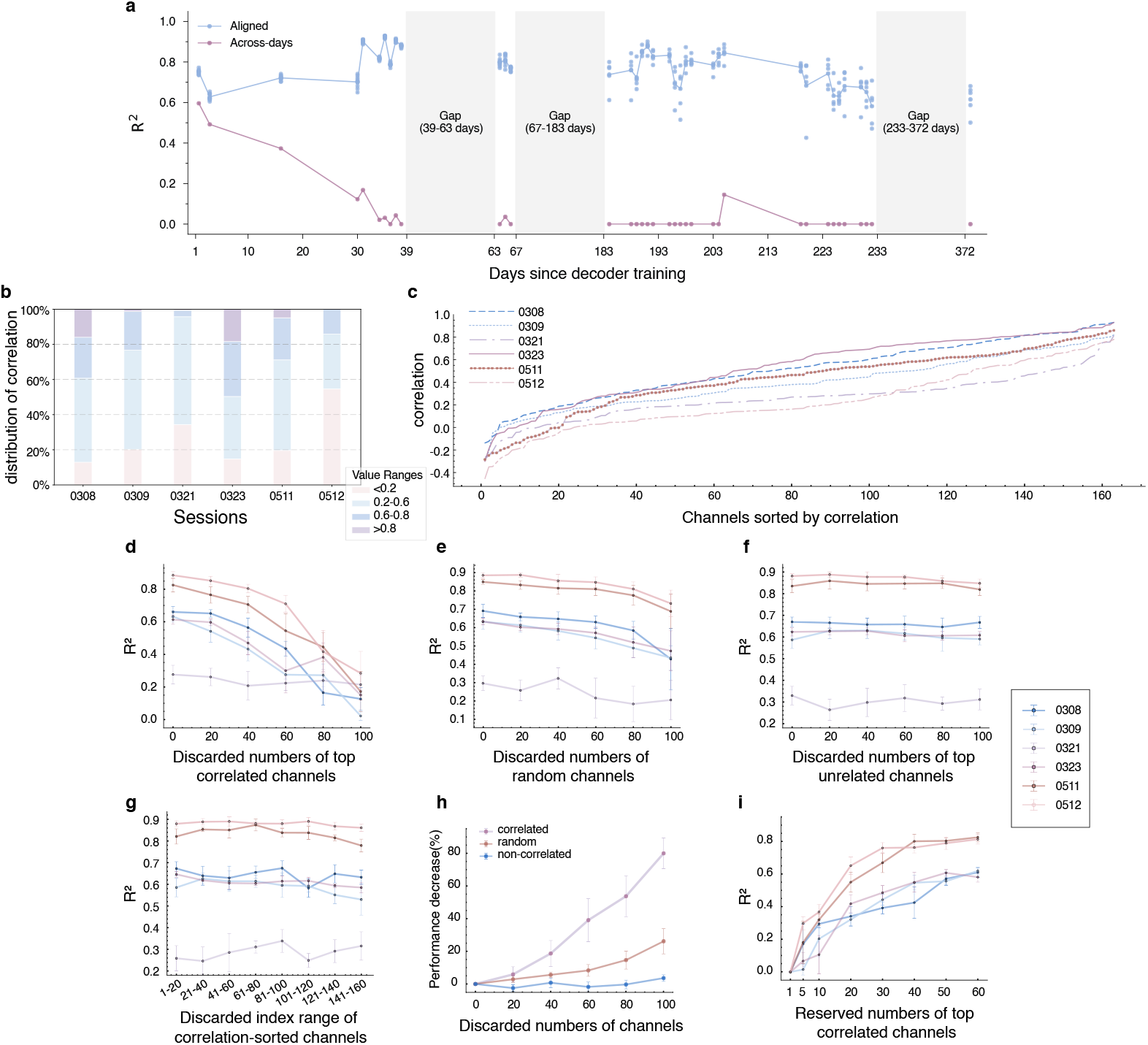
Small number of stable multiunits is enough for decoding across days. **a**, Behavior prediction performance using a fixed decoder. Each dot represents performance in one individual experiment. Claret line quantifies the progressive performance degradation of a fixed decoder under conditions lacking alignment interventions. The performance of a fixed decoder after alignment with meta aligner is shown for indigo. **b**, shows the distribution histogram of the correlations (corresponding to **c**). **c**, Sorted correlations between threshold crossing rates of identified sessions used for fixed decoder training and that of latter sessions. Specifically, trials of sessions are aligned to movement onset and computed averaged rates of trials which sharing same target direction in all 163 identified channels independently. For every channel, computing its correlations of 8 targets with corresponding trial-averaged rates respectively, and the mean of 8-targets correlations represents every channel-pair correlation. For comparison, **d, e** and **f** denote individual experiments that evaluating decoding performance of Meta-AlignNN in three conditions respectively. **d** shows decoding performance as a function of numbers of discarded top correlated channels (every channel and its correlation is showed in **c**; we set rates of channel at every time step to be zero in practice for “discarding” the channel). Error bars represent s.d. for 6 individual experiments. Same with **d**, except that **e** ignored random channels and **f** ignored top non-correlated channels. **g**, decoding performance as a function of different range of discarded correlation-sorted channels. **h** illustrated the relationship between decoding performance decrease and the number of discarded channels in three conditions, which is corresponding to **d, e** and **f** respectively. Error bars represent s.d. for each session. **i** shows how performance is with numbers of channels.

Given its impressive performance, we are curious how Meta-AlignNN maintains stability when dealing with neural activity instability over a span of more than a year. Figure 5c shows the sorted correlations between threshold crossing rates of identified sessions used for fixed decoder training and that of latter sessions. Channels that holds correlation from 0 to 0.6 is the biggest part of all 163 channels although proportion varies in different sessions. Significantly, in session 20230321, only about 10 channels exhibit correlation values above 0.4, which helps explain the poorer performance observed in Figures 5d,e,f,g for that session. Figure 5b further displays a histogram distribution corresponding to Figure 5c. Apart from 20230321, other sessions show a consistent pattern. Figure 5g prove that discarding different range of same 20-length channels has little impact on decoding and the discarded range with higher correlations tend to show worse performance. The decoding performance decrease severely and slightly in Figures 5d,e, respectively, while almost no decline in Figure 5f. The average decrease of all sessions except 20230321 is shown in Figure 5h.

From these findings, we draw an important conclusion: Meta-AlignNN leverage information contained in limited stable and correlated channels to help alignment and stabilization, which is similar with (Degenhart et al., 2020). However, there is a key distinction. For (Degenhart et al., 2020), a channel selection algorithm is needed to explicitly identify channels for subsequent alignment; while for Meta-AlignNN, we do not explicitly constrain the model to use specific channels. Instead, the meta aligner learned to extract the useful knowledge and channels implicitly with meta training. Reasonably, the meta aligner pay more attention to high correlated channels spontaneously for the purpose of quick adaptation and alignment. Last but not least, Figures 5d,h show that decoding performance only decrease about 20% even top 40 correlated channels are ignored, which means that Meta-AlignNN is robust and insensitive with the number of high correlated channels, i.e., a few high correlated channels are enough for alignment and stabilization. More intuitively, Figure 5i shows that Meta-

AlignNN with 20 channels already get not bad result and nearly saturated performance can be achieved in 40 channels. This finding is beneficial for the promotion and application of BCI, as Meta-AlignNN enables BCI stability with only a small number of channels, significantly reducing the reliance on high-throughput recordings.

### Meta-AlignNN achieves robust cross-subject alignment and scalable cross-task generalization via behaviorally anchored latent dynamics alignment

Our results demonstrate that Meta-AlignNN can effectively capture latent dynamics preserved across subjects and tasks underlying similar behaviors, thereby exhibiting robustness and scalability for generalization across different task paradigms and subjects.

As illustrated in Figure 4d, despite the complex diversity of activity patterns across subjects, the similarity between latent trajectories in Figure 4d2 and Figure 4d3 proves that Meta-AlignNN can capture preserved latent dynamics across subjects. As previously explained, “RS -> NW” in Figure 6 means that Monkey_RS uses the js-8 decoding model from Monkey_-NW to decode its own js-8. Figure 6a explained more clearly that Meta-AlignNN align the latent dynamics effectively across subjects from a session pair perspective (see Methods). Figure 6c calculates the behavioral correlation and latent variable correlation across session pairs with 12 test sessions (see Methods). However, due to the monkeys being well-trained, the behavioral correlation for session pairs is high and shows no significant differences. Therefore, Figure 6d analyzes this issue at a finer granularity, focusing on trial pairs (for clarity, only trial pairs with behavioral correlation less than 0.8 are included; see Methods). We find that as the behavioral correlation increases, the latent variable correlation is more likely to be higher. Figure 6e quantitatively describes the relationship between behavioral correlation and latent variable correlation, indicating that shared behavioral constraints drive preserved dynamics across subjects, consistent with the findings of (Safaie et al., 2023). This preservation facilitates cross-subject decoding performance comparable to within-subject decoding, as shown in Figure 6b.

**Figure 6.**
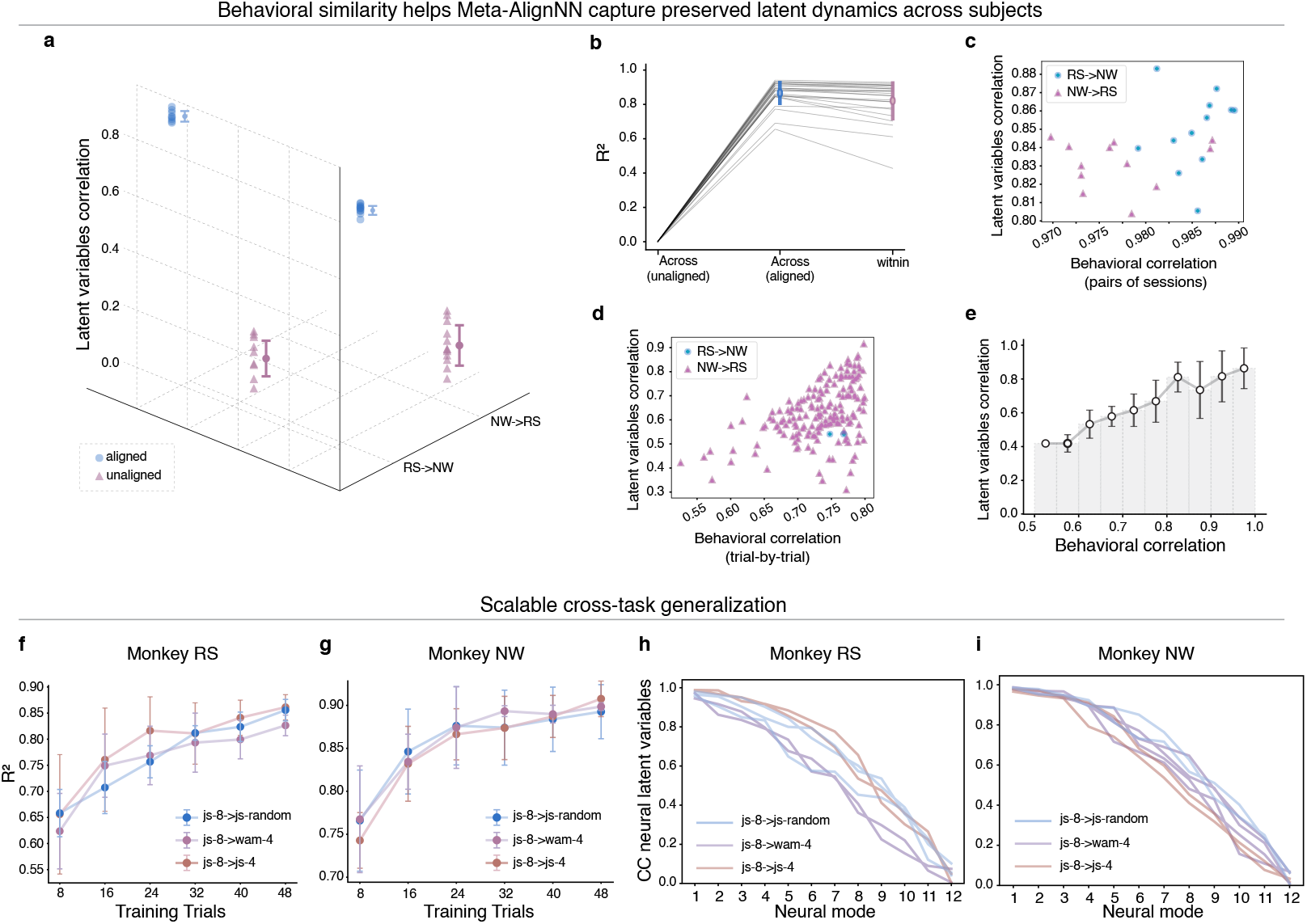
Meta-AlignNN achieves scalable cross-subject and cross-task generalization by aligning behaviorally anchored latent dynamics. **a**, represents the latent variables correlations of all trials averaged over the eight target directions. Each dot is the correlation of session pair. **b**, Decoder trained on sessions from one monkey is employed to predict the behavior of another monkey with alignment (indigo). These outcomes are compared to within-session condition (claret; decoders trained and tested within the same session) as well as those without alignment. Each line represents an individual comparison for each session across monkeys. **c** and **d**, illustrate that the latent dynamics between monkeys are linked to the similarity in their behavioral patterns. **c**, For each session pair, and **d**, for each trial pair with behavioral correlation lower than 0.8, single dots are color-coded by alignment order. **e**, Latent variables correlation as a function of behavioral correlation. Error bars represent s.d. for all trial pairs. **f** and **g**, When using Meta-AlignNN to align js-8 with different tasks (js-random, wam-4, js-4), the type of task has no significant effect on decoding performance on the test set with different calibration trials, in both monkeys. **h** and **i**, Similarly, the CCs for neural latent variables evaluated on test set show no significant differences across different cross-task types, in both monkeys.

Furthermore, the principle extends effectively to cross-task scenarios. Despite differences in task paradigms (joystick control vs. touchscreen reaching) and target numbers (8-target vs. 4-target vs. random-target), Meta-AlignNN achieved Comparable decoding performance (Figure 6f,g) and CCs of neural latent variables (Figure 6h,i) across all task combinations (js-8 -> [js-random, js-4, wam-4]). This scalability also stems from behaviorally anchored latent dynamics, specifically: 1. In js-random, js-4, and js-8, the hand movements controlling the joystick and the cursor movement directions are consistent; 2. While wam-4 involves touchscreen reaching and js-8 involves joystick control, the hand movement directions and target positions are still aligned (e.g., “reaching upward to hit a *whack-amole* target appearing at the top of the screen” corresponds to “pushing the joystick upward to move the cursor to a target at the top of the screen”). The behavioral invariance enables Meta-AlignNN to disregard superficial task differences while focusing on preserved movement-related dynamics. This scalable cross-task generalization has profound implications: clinicians could collect foundational data from simple tasks, then flexibly adapt decoders to new tasks through behaviorally anchored latent dynamics alignment.

The above results demonstrate Meta-AlignNN’s dual-alignment capability: 1) Within cross-subject alignment, it leverages natural behavioral correlations between subjects to identify shared neural manifolds; 2) Within cross-task alignment, it utilizes behavioral equivalences across different task implementations to extract invariant control signals. This intrinsic unified mechanism forms the foundation of Meta-AlignNN as a unified BCI decoding framework across time, tasks, and subjects, endowing it with strong robustness and scalability. This has important implications for the development of clinical BCIs: it enables “plug-and-play” adaptation, allowing new individuals or tasks to be integrated into the existing BCI system with only a subset of trials, thereby meeting the clinical demands for long-term, efficient, and stable usability across patients and tasks.

### Real-time brain-control

Meta-AlignNN achieved significant success in various across-time, across-task, and across-subject experiments (offline decoding and real-time brain-control) conducted over nearly two years involving three monkeys (Monkey_RS, Monkey_NW, Monkey_HF) and four tasks (js-8, js-random, *whack-a-mole*, and *Black Myth: Wukong*). **Supplementary Video 1** demonstrates realtime brain-control examples in each generalization setting. Left: Monkey_NW successfully performed the js-8 task with Meta-AlignNN’s assistance, 100 days after the decoder was trained (across-time). Center: Monkey_-NW used a decoder trained on Monkey_RS to perform js-8 efficiently with Meta-AlignNN (cross-subject). Right: Monkey_RS performed js-8 accurately using a decoder trained on the wam-4 task, aided by Meta-AlignNN (across-task). It is important to note that in these experiments, the monkeys’ hands were fixed, meaning they could not perform actual hand movements, with decoded behavior served as its visual feedback. As a result, real-time brain-control force the model to decode the behavior from its ideas and intentions precisely, otherwise terrible prediction will make the monkey tired and fretful through visual feedback, which then reduce the willingness and motivation of monkey to complete the task. It will conversely influence decoding performance, leading to a vicious circle. Undoubtedly, Meta-AlignNN performs well in real-time brain-control, demonstrating excellent performance across time, subjects, and tasks (Figure 7), evaluated by trial completion efficiency. Additional longer-duration brain-control results are shown in Figure S9. Due to poor brain-control performance, both fine-tuning and GRU led to a sharp decline in the monkey’s engagement, so only a few minutes were recorded (after which the experiment was discontinued). In contrast, Meta-AlignNN consistently maintained stable performance.

**Figure 7.**
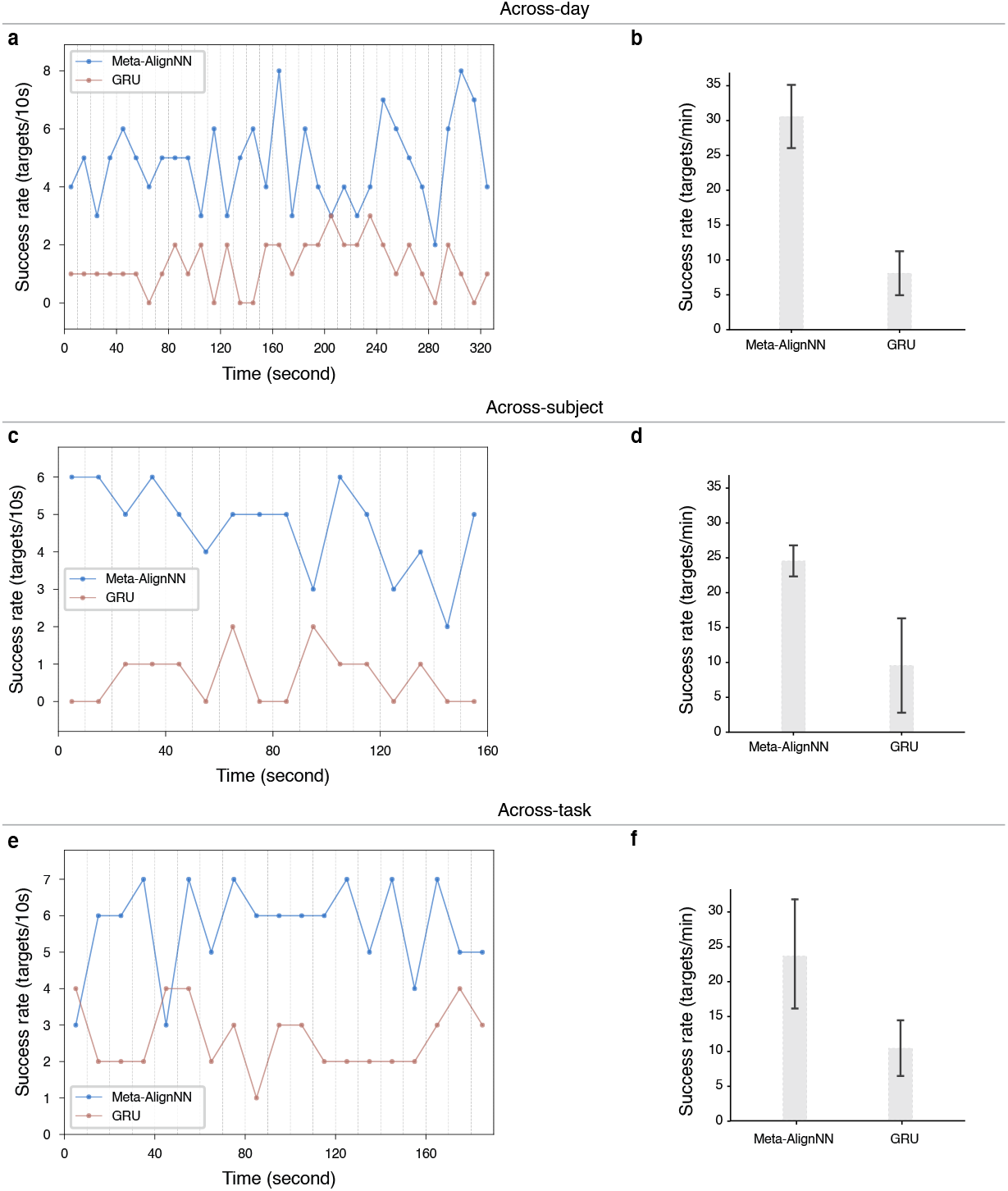
Real-time brain-control results across time, subjects, and tasks. **a**, Comparison of brain-control results between Meta-AlignNN and GRU in a specific testing session under the across-day scenario, measured by the number of targets hit every 10 seconds in js-8. **b**, Statistics of brain-control results between Meta-AlignNN and GRU across multiple testing sessions in the across-day scenario, measured by the number of targets hit per minute in js-8. Error bars represent s.d. for all test sessions. Similarly, **c** and **d** respectively show the brain-control performance in the across-subject scenario (MK_RS <-> MK_NW), while **e** and **f** respectively show the brain-control performance in the across-task scenario (js-8 -> wam-4).

To further evaluate Meta-AlignNN’s ability to align low-dimensional neural dynamics across time, individuals, and tasks simultaneously, we designed a novel experiment. First, we constructed a decoding algorithm using neural activity recorded in July 2023 from the Monkey_RS while playing wam-4. Then, with the help of Meta-AlignNN, we were pleasantly surprised to find that Monkey_HF could seamlessly control the game *Black Myth: Wukong* in January 2025 using this decoding algorithm from a different time, individual, and task (Figure S10). For details on across-task setting between wam-4 and *Black Myth: Wukong*, please refer to **Across-task paradigm design** method. In *Black Myth: Wukong*, completing a specific level (defeating two minions and one boss) is referred to as one round. The number of rounds Monkey_HF can complete per hour, evaluated with several sessions, is shown in Figure S10b. With only 5 minutes of fine-tuning data, Meta-AlignNN outperformed and was more stable than the GRU trained with approximately 40 minutes of data collected on the same day while Monkey_HF played *Black Myth: Wukong*. In contrast, a decoder model trained on data from the past 20 days of Monkey_HF playing *Black Myth: Wukong*, fine-tuned with 5 minutes of data from the same day, failed to perform effectively. More intuitively, Figure S10a shows a comparison of the rounds completion between these three models within a single hour. As demonstrated in **Supplementary Video 2**, Monkey_HF was using Meta-AlignNN to play *Black Myth: Wukong* across time, subjects, and tasks.

## Discussion

In this study, we present a unified meta-learning framework, Meta-AlignNN, designed to address the limitations of previous approaches that focus solely on BCI stability along a single dimension (e.g., time), and to ensure BCI stability and robustness across subjects, time, and tasks. Extensive experiments and evaluations on public datasets demonstrated that Meta-AlignNN outperformed FA stabilization (Degenhart et al., 2020), ADAN (Farshchian et al., 2018), SABLE (Jude et al., 2022), and DANN (Ganin et al., 2016) in decoding accuracy and data efficiency. Additionally, Meta-AlignNN successfully supported real-time brain-control in *Black Myth: Wukong*, illustrating its capability in simultaneous across-time, across-subject, and across-task alignment of neural dynamics.

Meta-AlignNN exploits the intrinsic behaviorally anchored low-dimensional structure of neural population activity, and aligns well with evidence that movement-related neural manifolds are preserved despite electrode instabilities (Pandarinath et al., 2018; Gallego et al., 2020), as well as the theoretical foundation of stable neural-behavior mappings (Chestek et al., 2007; Stevenson et al., 2011; Flint et al., 2016). Unlike existing approaches using autoencoders (Pandarinath et al., 2018), adversarial training (Farshchian et al., 2018; Jude et al., 2022), or linear methods (Degenhart et al., 2020; Gallego et al., 2020), our approach addresses the fundamental challenge of dynamically mapping non-stationary neural signals to stable, behaviorally interpretable latent dynamics by using the meta aligner to infer the optimal encoding function. Our results and analyses align closely with recent studies emphasizing neural pattern conservation across subjects, time, and tasks (Gallego et al., 2018, 2020; Safaie et al., 2023). Meta-AlignNN can effectively capture preserved latent dynamics underlying natural behavioral consistency, thereby exhibiting robust and scalable generalization across long timescales, diverse task paradigms, and subjects. Additionally, we found that Meta-AlignNN can achieve BCI stability with only a small number of stable recording channels, significantly reducing the reliance on high-throughput recordings.

Three key considerations for implementing Meta-AlignNN in BCIs are as follows. 1. Electrode/session selection: Our method accommodates existing electrode/session selection protocols (Degenhart et al., 2020; Fraser and Schwartz, 2012; Dickey et al., 2009; Tolias et al., 2007), with flexibility to incorporate custom criteria for identifying stable neural signals. 2. Meta-alignNN flexibility: The framework supports cross-time, cross-subject, and cross-task adaptation through task definition and data collection adjustments while retaining its core architecture and procedure. 3. Decoder compatibility: Since Meta-AlignNN imposes no theoretical constraints, it can be combined with any decoder architecture, such as Kalman filter (Wu et al., 2006; Wu and Hatsopoulos, 2008; Gilja et al., 2012), Naive Bayes (Barbieri et al., 2005; Kloosterman et al., 2014), Wiener filter (Carmena et al., 2003), Wiener cascade (Pohlmeyer et al., 2007), Support vector regression (Smola and Schölkopf, 2004; Chang and Lin, 2011), XGBoost (Natekin and Knoll, 2013), Feedforward neural network (Orbach, 1962; Goodfellow et al., 2016), or Simple RNN (Rumelhart et al., 1986; Goodfellow et al., 2016).

A notable limitation is Meta-AlignNN’s reliance on substantial training sessions to capture variability effectively. On the one hand, generating synthetic neural data aligned with natural recording distributions, potentially via methods (Degenhart et al., 2020) or advanced AIGC techniques, represents a promising direction to address this issue. On the other hand, as large-scale neural datasets continue to emerge, Meta-AlignNN shows exceptional scalability potential and becomes increasingly powerful.

In future work, the versatility of Meta-AlignNN could be further explored in more complex and varied tasks, as well as in larger-scale and more diverse subject populations.

Our findings establish a unified solution for maintaining BCI stability, robustness, and scalability across subjects, time, and tasks, providing the foundation for meeting the clinical demands of long-term, efficient, and stable usability across patients and tasks.

## Methods

### Behavioral task

Overall, we trained three rhesus macaques (Monkey_-RS, Monkey_NW, and Monkey_HF) to perform four different tasks (center-out reaching: referred to as “js-8”, random-target reaching: referred to as “js-random”, *whack-a-mole*: referred to as “wam-4”, and *Black Myth: Wukong*). We trained Monkey_RS and Monkey_NW to sit on a primate chair and used a custom flat joystick to play js-8 and js-random. In addition, we also trained them to play the touchscreen *whack-a-mole* game (see Figure 1a). Over the course of two years, we collected data from more than 300 sessions, with session intervals ranging from 1 day to 30 days. In js-8, the cursor was reset to the center of the screen at the beginning of each trial. Following a randomized delay interval, the monkey was cued to execute reaches toward one of eight radially arranged peripheral targets (angular spacing: 45°), with target locations pseudorandomly selected with uniform probability. Successful trials requiring target acquisition within a 3-second temporal window triggered liquid reward delivery. Earlier, the monkeys were using a planar manipulandum to control the cursor, while later a robotic arm was controlled instead where the fingertip of robotic arm moved on the plane that parallel to the screen and 5 millimeters away. The 2D position is acquired by projecting 3D position of the fingertip to the screen which equals the positions of the robotic arm moving plane. The moving velocities of cursor and robotic arm with respect to planar manipulandum are the same. However, robotic arm will hit the target with fingertip along the vertical direction of the screen if its 2D position reach the target range, which is extra action compared with cursor control. Throughout task execution, continuous kinematic tracking of the effector (cursor/robotic arm) was performed at 20 kHz resolution, with synchronized digital event markers precisely timestamping behavioral transitions across trial phases (e.g., stimulus onset). Similar to js-8, but in js-random, the targets are random, and at the beginning of each trial, the cursor’s starting position is immediately after the previous trial (instead of being initialized to the center of the screen). In wam-4, at the start of each trial, four gopher holes appear on the screen (top, bottom, left, right). A gopher then randomly appears in one of the holes. We train Monkey_RS and Monkey_NW to hit the gopher by touching the screen within a specified time, and correct hits are rewarded with liquid food. Monkey_HF was trained to play *Black Myth: Wukong*. We selected a segment of the game (which involves defeating two minions and one boss) as the training content for the monkey. The monkey needs to control the joystick to move the character forward and press buttons to make the character attack. When Monkey_HF moves the character and defeats the boss to complete the game level, the character will return to the starting point of the level and repeat the process.

### Neural implants

To obtain chronic electrophysiological recordings from cortical neuronal populations, we implanted eight self-developed 128-channel flexible micro-electrode arrays in the primary motor cortex (M1) and the dorsal premotor cortex (PMd) of monkeys, respectively, using standard surgical procedures to achieve 1024-channel (8×128) neural recordings. All experimental protocols involved in this study were approved by the Animal Management Committee of Lingang Laboratory (NZXSP-2022-1) and conducted in accordance with all relevant ethical regulations.

The Flexible micro-electrode arrays were fabricated using standard microfabrication techniques. Each flexible micro-electrode array consists of three 1-μm-thick polyimide (PI) layers alternating with two gold (Au) conducting layers in a sandwich configuration. This results in a total thickness of approximately 3 μm. The flexible micro-electrode array adopts a spiral design to ensure convenient implantation. The flexible microelectrode array consists of eight 20-μm-wide spiral filaments (shanks) and each filament incorporates an implantation hole at its tip. The recording sites on each filament were arranged with a 90-μm pitch over a total length of 1500 μm. Tungsten wires with sharp tip-to-base transition were fabricated through a controlled electrochemical etching process. The base diameter of the tungsten wires is 100 μm, whereas the diameter of the tip is about 30 μm. The filament sections of the flexible micro-electrode array were released from the substrate. The implantation hole of each filament was aligned and assembled with a tungsten needle during the implantation process. The released device was stored under sterile conditions.

Each electrode-array consisted of 8 shanks. Neural signals were acquired during behavioral tasks using the TRODES system (https://spikegadgets.com/trodes/), a available neurophysiological acquisition platform. Raw signals underwent digitization followed by bandpass filtering, and subsequently converted into multiunit activity via threshold crossing detection, with the threshold set to 22 μV based on waveform signal-to-noise characteristics. In this study, we focused on multiunit threshold crossings per channel to prioritize realtime responsiveness for online BCI control, avoiding the time-consuming process of offline single-neuron spike sorting.

### The Meta-AlignNN

The Meta-AlignNN model consists of two components, a meta aligner followed by a decoder. Specifically, the training process of Meta-AlignNN can be concluded as two-stage: (1) training a decoder with identified sessions and electrodes; and (2) training a meta aligner with abundant later sessions using optimization-based meta learning algorithm.

#### Identify sessions and electrodes for decoder

As mentioned above, the sessions and electrodes used for decoder training is one crucial aspect of Meta-AlignNN because the base latent space and its relationship with behavior learned by decoder would affect performance of Meta-AlignNN a lot. In our case, we manually select many consecutive days of recording sessions (one session per day) in which the monkey had good behavior, which means that the right column session in Figure S8a is preferred. Then we select the electrodes of each session during consecutive days that hold high signal to noise ratio, as shown in Figure S8b, the first two rows denoted good selected channels while the third row was bad discarded channels. Each subplot in Figure S8b contains a large number of spikes and green or red line denotes one random spike. Two horizontal lines represent threshold for detection. Notably, the last subplot in third row holds few recordings as corresponding electrode corrupted. Finally, we take union of channels selected in each session as the overall channels for both decoder and meta aligner training.

#### The decoder

The decoder can’t be called “decoder” in a strict sense because it consists of a GRU encoder followed by a regressor. We refer these two components as decoder just for convenience. Once sessions and electrodes were identified, we then used sliding window of length *T* to cut out the neural activities of successful trials in selected sessions and electrodes and the corresponding behaviors. The generative forward process of the decoder is as follows:

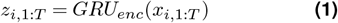

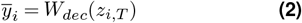

where *GRU*_*enc*_, *W*_*dec*_ are the parameters of the GRU used to encode gaussian smoothed multiunit threshold crossings rate into latent variables, and the parameters of the non-linear layers to decode behavior from the latent variables respectively. Significantly, we do compute the last prediction of output sequence, i.e., 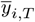. For simplicity, we denote it as 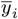, where Eq. 2 shows.

The Adam optimization algorithm (Kingma and Ba, 2014) is applied to minimize the defined loss function at every training iteration:

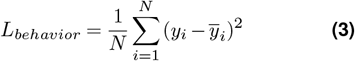

where *y*_*i*_ is the true behavior of one sample, 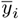 is the predicted behavior, and N is batch size.

#### The meta aligner

Once the decoder was well trained with the former identified sessions and electrodes, it means that a proper latent space was extracted for neural BCI decoding. However, due to the variation in neural recordings across time, subjects, and tasks, the decoder may not be usable for future decoding. As mentioned above, the mapping between latent neural dynamics and motor intent exhibits temporal stability (Chestek et al., 2007; Stevenson et al., 2011), one critical solution for long-term BCI decoding is thus to find the powerful aligning function to project nonstationary neural signals into the stable latent subspace, thereby enabling reliable behavioral inference. The meta aligner can be viewed as this powerful aligning function. We select transformer encoder as the architecture of meta aligner considering its powerful ability of feature extraction and representation (Vaswani et al., 2017; Devlin et al., 2018).

#### Training procedure

Inspired by meta learning, we incorporate the training procedure and policy of MAML (Finn et al., 2017) into our Meta-AlignNN framework to train the meta aligner. The core objective of this training paradigm is to develop a meta aligner capable of fewshot adaptation to novel recording sessions with neural signal shifts, utilizing minimal neural data and computational iterations. To achieve this, the meta aligner undergoes meta-training across a heterogeneous task distribution, enabling rapid generalization to unseen tasks across time, subject, and task with sparse data samples. The meta aligner, denoted as *ψ*, is defined to align the neural activity. The meta aligner is conditioned during the meta-training phase to achieve generalization through exposure to diverse task distributions. Different from the meaning of “task” in across-task decoding (referring to behavioral tasks), here the meaning of the term “task” (in the context of meta-learning) is that feeding training data of one latter session recording as the input of meta aligner and using its output aligned data to predict the behavior with the fixed decoder. Formally, each task *T* = {*L*(*x, y*), *s*(*x, y*), *q*(*x, y*)} consists of a loss function *L*, training data *s*(*x, y*), and testing data *q*(*x, y*). The loss function here is actually Eq. 3. In real implementations, 80 percent of recording sessions that spanning across several months are assigned to form training tasks, while 20 percent to form testing tasks. As shown in Figure S2c, to train meta aligner, the data were organized, partitioned, and collected in slightly different ways across three application scenarios: across days, across subjects, and across behavioral tasks. Each task also contains training data and testing data. We denote the distribution over all tasks as *p*(*T*).

The meta aligner is mathematically formulated as a parametrized function *h*_*ψ*_, where *ψ* denotes its learnable parameters. For ease of explanation, we introduce the training procedure if only considering one training task. First, the meta aligner is trained with *K* samples sampled from *s*(*x, y*) of task *T*_*i*_ and its parameters *ψ* will become 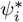 through gradient descent updates on corresponding loss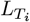. The number of gradient descent updates is unconstrained, thereby permitting the execution of one or more iterative optimization steps. To streamline mathematical notation, we adopt a single gradient descent step in all subsequent formulations and analyses presented in this work, and the formula is shown below:

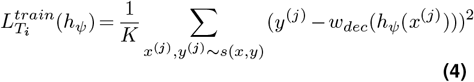

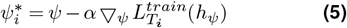

where *w*_*dec*_ is the parameters of the fixed decoder and learning rate *α* is a hyperparameter. Second, we need to test on new samples from *q*(*x, y*) of task *T*_*i*_, the meta aligner’s parameters *ψ* are optimized via the test error 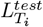 evaluated on data sampled from *q*(*x, y*), ensuring that the adapted parameters 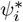achieve minimal test error:

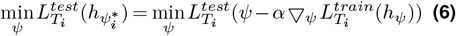

The training process of one training task is showed vividly in Figure 2c.

The meta-training loss is computed by aggregating the generalization errors across all tasks sampled from the task distribution *p*(*T*). Formally, the meta-optimization objective is defined as:

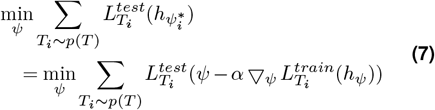

The meta-objective with respect to *ψ* is optimized via stochastic gradient descent:

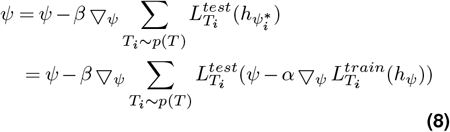

The meta-learning rate is denoted by *β*. Crucially, the meta-optimization operates on the meta aligner parameters *ψ*, while the meta-objective’s evaluation depends on the post-adaptation parameters 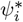, where each 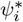 corresponds to a task-specific adaptation. As a result, gradient through gradient is needed when perform meta-optimization, which is computational expensively. As an alternative, we can use a first-order approximation to ignore second-order derivatives, which is proved to perform well on some benchmarks and provided with related theoretical analysis (Nichol et al., 2018).

As a conclusion, we consider each alignment as a task, and internal representations across all these different alignments are transferable and can be broadly applicable to new unseen alignment, which means meta aligner can make fast adaptation to unseen sessions.

### Across-task paradigm design

To validate the generalization of Meta-AlignNN in across-task scenarios, we designed several across-task combinations, including js-8 vs. js-4, js-8 vs. js-random, js-8 vs. wam-4, and wam-4 vs. *Black Myth: Wukong*. The specific task definitions and processing are shown in Figure S7. In detail, we processed the raw data of js-random, that is, pulling the starting point of each trial to the center rather than starting from the end of previous trial in real experiments. To compare with js-8, we divided the screen area into eight quadrants and aligned the starting point of each trial to the center of the screen. After center alignment, the trajectories were distributed across these quadrants. These eight quadrants are considered as the 8 directions of js-random, as shown in Figure S7a. There is no doubt that freedom of operation in js-random is much higher than that in js-8. In order to investigate the influence of task complexity in aligning across tasks, we manually removed the 4 directions in js-8, namely js-4. Js-8 and js-4 are another set of experiments used across tasks. (Gallego et al., 2018) claims that flexibly combined, well-preserved neural modes may underlie the ability of M1 to learn and generate a wide-ranging behavioral repertoire, and at the neural population level, the structure and activity of the neural modes is largely preserved across tasks. Based on this, we are curious about whether across-task alignment of Meta-AlignNN in different context is still effective. We designed a set of tasks for wam-4 and js-8. The monkey uses the manipulandum to control the cursor to reach 8 targets in js-8, while it directly touches the 4 targets of the screen with hands in wam-4. When decoding, the former usually uses regression to decode kinematics, while the latter usually uses classification to decode intention. We were interested in whether alignment across tasks with Meta-AlignNN still work in these two different contexts. Here, We have done some special processing for data and decoding in wam-4. First, we convert the behavior of hitting the target into a “fixed trajectory on the screen” through affine transformation (Figure S7b). Of course, the trajectory here is imaginary and only serves as the ground truth for decoding. Each hitting target position corresponds to a fixed trajectory. The reason for this operation is that we believe that the neuron population activity patterns of direct arm strikes and manipulandum operations are similar when the behaviors are similar (for example, they are both in the upward direction). Secondly, after decoding the trajectory, we will map it back to the specific target position according to the direction of the trajectory. The decoding method of wam-4 has been changed from classification to regression. In this way, wam-4 can be decoded using the decoder trained with js-8 training data and js-8 vs. wam-4 can be the third set of experiments used across tasks. In the across-task comparison between wam-4 and *Black Myth: Wukong*, wam-4 involves five classes (hitting the four directional gopher holes and doing nothing), while *Black Myth: Wukong* involves three classes (move forward, attack, and do nothing). Considering the behavioral similarity, the following mappings are made: “Reaching to hit the ‘upper’ gopher” (wam-4) corresponds to “moving forward” by pushing the joystick up in *Black Myth: Wukong*; 2. “Reaching to hit the ‘right’ gopher” (wam-4) corresponds to “attacking” by pressing the button on the right side of the joystick in *Black Myth: Wukong*; 3. “Doing nothing” (wam-4) corresponds to “doing nothing” in *Black Myth: Wukong*. With this setup, wam-4 decoding model is used to control *Black Myth: Wukong*.

### FA stabilization

We adopted the same procedure that (Degenhart et al., 2020) mentioned, including 1. training initial baseline stabilizer and decoder; 2. using three computational phases to update the stabilizer: recomputing factor analysis parameters, identifying stable electrodes and aligning the coordinate system for the neural manifold across days; 3. decoding movement trajectories using initial decoder with aligned neural activity except that we replaced kalman filter with GRU as decoder. To obtain the evaluation metrics shown in Table 1, such as the mean R-squared for 20230511. We first trained the initial stabilizer and decoder using trials from 20230510, then updated the stabilizer with early trials from 20230511, and finally used decoder to evaluate the performance with remaining stabilized trials in 20230511.

### ADAN

We followed the official implementation of (Farshchian et al., 2018) to obtain the evaluation metric in Table.1 For example, to calculate the mean R-squared for 20230511, firstly, we used session 20230510 for training an initial within-day model, which integrates an autoencoder module with a downstream predictive network. Then, we aligned neural activities of session 20230511 to those of 20230510 with the optimization of ADAN architecture. Consequently, aligned latent representations of neural activities were translated into behavior movement using the previous trained predictor.

### DANN

We consider DANN as one comparison model due to its ability to generate domain-invariant features. Here, each session is regarded as each different domain, and domain-invariant features that are insensitive to change of neural activity can then be used for BCI decoding. The implementation for the result achieved by DANN that is shown in Table 1 almost followed the methods mentioned in the original DANN paper (Ganin et al., 2016). We defined test session and its adjacent former session as target domain and source domain, respectively, e.g., 20230510 as source domain and 20230511 as target domain when testing on 20230511. Next, we changed the classifier in DANN as GRU regressor and followed the same domain-adversarial training method. The evaluation performance is shown in Table 1.

### SABLE

For comparative analysis, we replicated the SABLE architecture (Jude et al., 2022), which integrates a sequential variational autoencoder with a decoder for behavior prediction. Crucially, SABLE employs a gradient reversal layer connecting the neural decoder and encoder, adversarially training the model to encode session-invariant latent dynamics necessary for robust behavioral decoding. To obtain the performance listed in Table 1, we trained SABLE on five consecutive sessions together with part of a subsequent held out session and test on the rest of that session.

### Neural latent trajectory and latent space analysis

To get 2D t-SNE visualization of Meta-AlignNN’s latent space showed in Figure S5, the detailed implementation can be summarized as below. Firstly, only successful trials were included for analysis. All trials were aligned to their end, and a fixed time window ending at the trial’s completion is extracted. Under this way, a large number of trials were obtained from both the consecutive sessions used to train the decoder and the test sessions (in this case, 20230511 and 20230512). Secondly, neural recording of these trials are projected into latent representations in three ways (within, aligned, and unaligned). Here we use the output of GRU encoder (i.e., the result of Eq. 1, which is a part of decoder) as latent representations, with each trial corresponding to one latent representation. For comparison, trials extracted from consecutive sessions were directly fed into the decoder to obtain their latent representations, while trials from test session were used in a controlled experiment (i.e., aligned with the meta aligner or not). Finally, the aligned or unaligned outputs were fed into the decoder to obtain their corresponding latent representations. For visualization and analysis, the three types of latent representations mentioned above were reduced to two dimensions using t-SNE, with different colors and markers used to indicate different trial target directions and representation types.

Figures 4a,d,g,j,m show the two-dimensional t-SNE visualization of mean neural latent trajectories for sessions used for training decoder and tested sessions before or after alignment with Meta-AlignNN. The implementation details are as follows. Unlike the trial extraction method mentioned earlier in Figure S5, here all trials were aligned to the movement onset, and it is not required for trials to have the same length. The neural activity of each trial was binned at 40 ms and preprocessed. A sliding window (moving one bin at a time until the end of the trial) was used to segment the neural activity, which was then fed into the decoder using three ways (within, aligned, and unaligned) like what mentioned above in Figure S5 and converted into latent representations. Then each trial was denoted as one latent representation sequence. Averaging the latent representation sequences of trials with the same target direction helps obtain a latent representation sequence corresponding to the behavior of reaching a specific target direction. Finally, by applying dimensionality reduction to the latent representations, latent neural trajectories can be visualized, where the trajectories themselves encode temporal information and the colors of the trajectories indicate target directions.

### Canonical correlation for neural latent variables

To investigate the improvements achieved by using Meta-AlignNN after alignment across time, tasks, and subjects, we applied Canonical Correlation Analysis (CCA) (Bach and Jordan, 2002) to compare the similarity between the aligned neural latent variables from the test session and the “within” neural latent variables. We calculated the canonical correlation of neural latent variables between sessions as follows: Following the method mentioned in **Neural latent trajectory and latent space analysis** section, each trial within a session was transformed into its corresponding latent representation sequence. Trials with the same target direction within each session were then sequentially concatenated. As a result, each session is represented as an ordered set of concatenated latent representation sequences for all trials. The canonical correlation between the neural latent variables of the session pair can then be calculated using the two latent variables matrices, *A* and *B*, formed from the sessions. Specifically, for each session, we project the neural latent variables onto the *m* principal component neural modes, resulting in two matrices, *L*_*A*_ and *L*_*B*_, each of size *m* × *T*, where *T* is the total time length of all concatenated trials in that session. CCA computes two linear transformations, for matrices *L*_*A*_ and *L*_*B*_ respectively, that project the data into new manifold directions in which the transformed representations from the two manifolds are maximally correlated.

Specifically, similar to the calculation process in (Gallego et al., 2018), the matrices *L*_*i*_ are first decomposed into orthonormal basis via QR factorization, 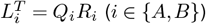, where the first *m* columns of *Q*_*A*_ and *Q*_*B*_ span the column spaces of 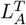*and* 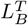, respectively. Next, computing the singular value decomposition for the constructed *m* × *m* inner product matrix of *Q*_*A*_ and *Q*_*B*_:

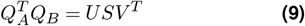

The diagonal entries of *S* are the pairwise canonical correlations for neural latent variables, listed from the largest to the smallest.

### Direction disorderliness of neural latent trajectories

To quantitatively describe the improvement in neural latent trajectories after alignment using Meta-AlignNN across time, tasks, and subjects, we proposed the metric “direction disorderliness.” Direction disorderliness refers to the angular deviation between the actual neural latent trajectories and the optimal neural latent trajectories, specifically the average angular deviation between trajectories with the same target direction. Here, the two-dimensional optimal neural latent trajectories correspond to straight lines in physical space from the center point to each target location (8 straight lines for js-8 and js-random, and 4 straight lines for wam-4 and js-4). In this calculation, the angle of a single trajectory is the mean of the angles of all the segments formed by connecting consecutive trajectory points. Additionally, the angles of trajectories with different target directions are anchored to one direction (by subtracting the angle of that direction’s trajectory), thus eliminating the rotational effect, to calculate the angular deviation between trajectories with the same target direction.

### Latent variables correlation and behavioral correlation

As described in the earlier subsection, the latent representation sequences of trials with the same target direction were averaged to obtain a representative latent representation sequence for each distinct target direction in the session. The Pearson correlation between the latent representation sequences corresponding to the same target direction in the two sessions was computed, and the average across all directions was taken to obtain the latent variable correlation for the session pair. Figure 6a shows the latent variables correlations computed using this method for 12 test sessions. When calculating the latent variable correlations between trial pairs (with same target direction) on a trial-by-trial basis, it refers to directly computing the Pearson correlation between the latent representation sequences of each trial pair. Similarly, when calculating the behavioral correlation between trial pairs on a trial-by-trial basis, it refers to directly computing the correlation between the cursor trajectories of the trial pairs. Figure 6d applies this method to all trial pairs across all test sessions and reports the results for those with behavioral correlation less than 0.8. Figure 6c calculates the behavioral correlation from the perspective of session pairs (12 test sessions), where the behavioral correlation for a session pair is the average of all trial-pair behavioral correlations within that session pair.

### Behavior decoding and real-time brain-control

We use Trodes to conduct multiunit threshold crossings detection on each recording channel, with threshold set to 22μV based on the signal-to-noise ratio of the waveforms. Subsequently, non-overlapping 40ms time bins were constructed from the threshold crossing counts on selected electrodes. For each recording channel, gaussian smoothing was applied to the binned counts to obtain smoothed threshold crossing rates. These were then used to generate inputs for Meta-AlignNN via a sliding window mechanism. Each neural input captured by a sliding window was fed into Meta-AlignNN for behavior prediction. The total number of inputs for one trial is related with the length and stride of sliding window. Sliding window mechanism is used for both online and offline behavior decoding, while sliding window ought to contain the latest neural activity in online mode.

## Acknowledgments

This work was supported by Lingang Laboratory (Grant No. LG-GG-202402-06), Shanghai Municipal Science and Technology Major Project (Grant No. 2021SHZDZX), the National Key R&D Program of China (Grant No. 2021ZD0203601 and 2019YFA0709504), National Natural Science Foundation of China (Grant No. 32221003 and 32161133024) and Shanghai Pilot Program (Grant No. JCYJ-SHFY 2022-010).

## Author contributions

Yongjie Zou was responsible for the design, training, and analysis of neural network models, algorithm coding, data analysis, figure generation, and manuscript writing. Zhengliang He was responsible for establishing the animal behavior experimental platform. Yongjie Zou, Kai Liu, Congcong Zhang, Fei Wang, Mingxin Li, Mengyu Li, and Yulei Chen jointly conducted the animal experiments. Yu Ling participated in part of the data analysis. Fei Wang, Mengyu Li, and Yulei Chen were responsible for the training of the animals. Yongjie Zou, Zhengliang He, and Chengyu T. Li participated in the discussion and revision of the manuscript. Shouliang Guan developed the flexible micro-electrode arrays. Shouliang Guan, Zhengliang He and Chengyu T. Li jointly supervised the study.

## Supplementary Information

**Figure S1.**
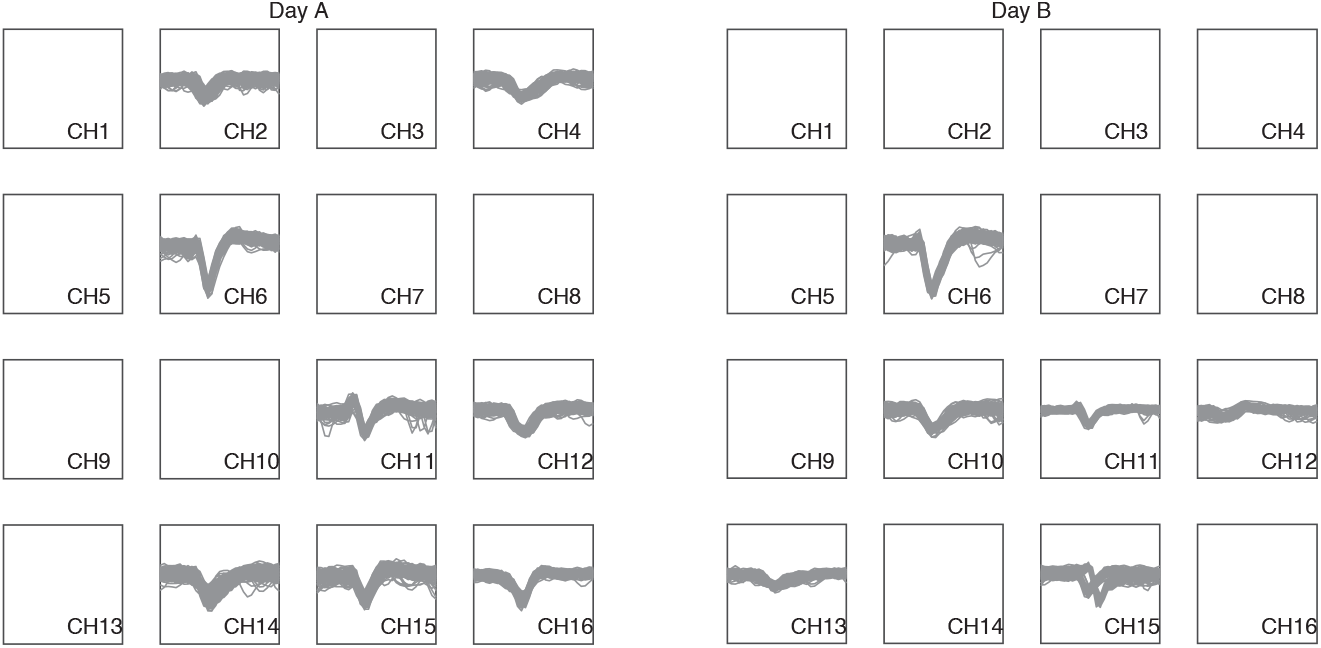
Signal recorded by electrodes across days. Multiunit threshold crossings for example recording channels on two different days for monkey RS, demonstrating substantial variations in signal recorded by electrodes across days.

**Figure S2.**
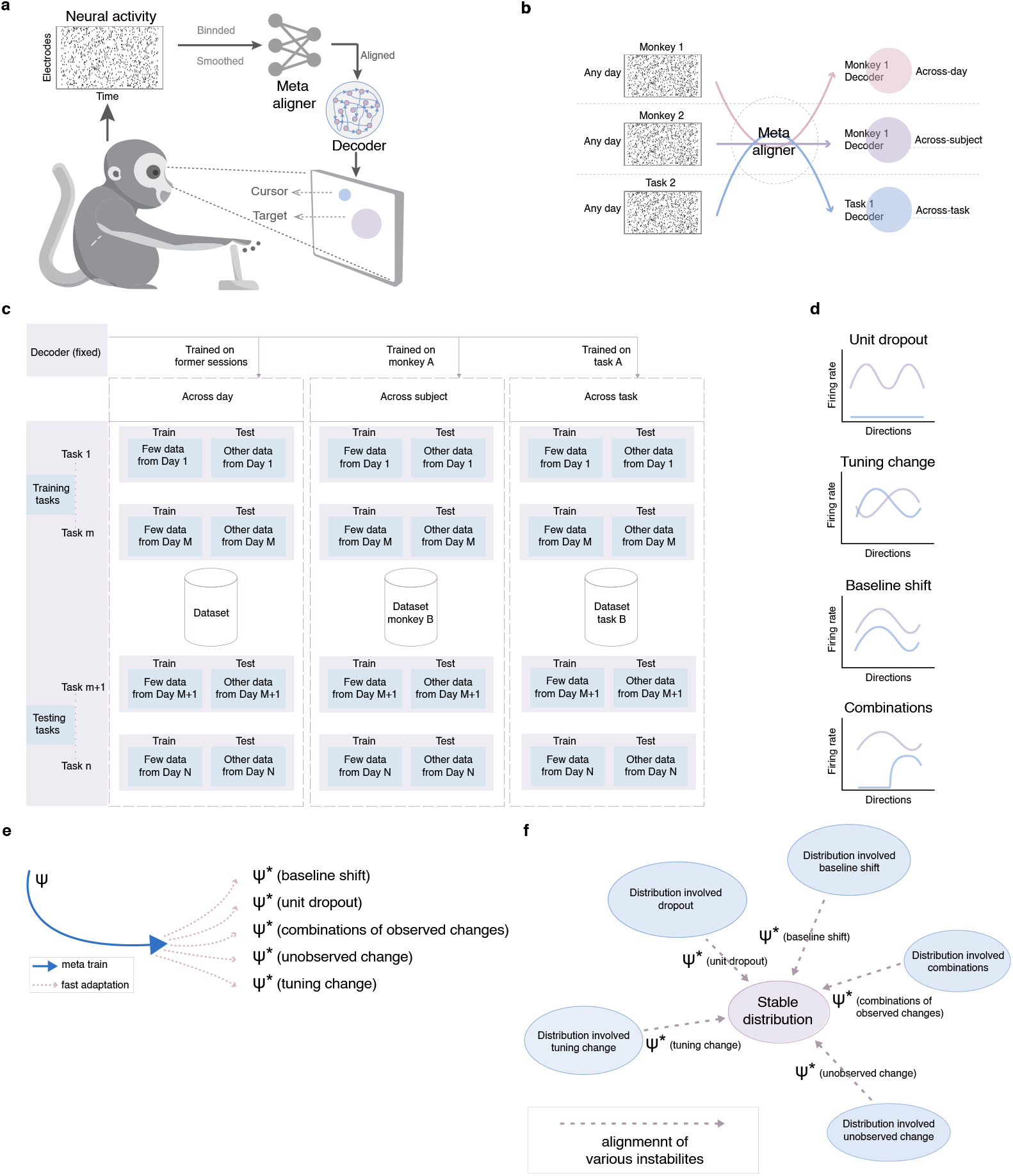
Supplementary details of Meta-AlignNN. **a**, Schematic of Meta-AlignNN-driven stabilization. Neural activity (visualized as spike raster representations) undergoes temporal binning and Gaussian smoothing before being processed by the meta aligner. The aligner transforms the input into aligned multiunit threshold crossing rates, which are subsequently decoded to estimate BCI cursor positions. Crucially, the meta aligner dynamically compensates for neural recording instabilities, while the decoder parameters remain fixed during operation. **b**, Integration framework of the meta aligner and decoder in three scenarios: across-day, across-subject, and across-task applications. **c**, Data partitioning protocol for Meta-AlignNN training in across-day/-subject/-task contexts. Training tasks are constructed using 80% of later recording sessions (one session/day, excluding earlier sessions used for decoder initialization), with the remaining 20% allocated as testing tasks. Each session (day) is treated as an individual task. **d**, Four simulated instability types applied to electrode subsets (defined by instability ratio): (1) Baseline Shift Instability: Introducing a fixed offset to threshold crossing rates in randomly selected electrode subsets; (2) Unit Dropout Instability: Nullifying threshold crossing rates in randomly chosen electrode channels; (3) Tuning Perturbation Instability: Swapping threshold crossing rates between equal-sized random channel pairs; (4) Composite Instability: Combined application of (1)-(3). **e**, From the perspective of parameter updating, the meta training policy of Meta-AlignNN implicitly enable the meta aligner quickly adapt to an unseen recording session with various instabilities. The blue solid line indicates the update process of *ψ* during meta training. Afterward, the meta aligner is able to reach various relatively optimal value rapidly (represented by red dashed line) for different corresponding alignment of instabilities. **f**, From the perspective of alignment in the data feature space, various instabilities are addressed with the alignment imposed by corresponding optimal meta aligner, i.e., *ψ*∗ in figure S2**e**.

**Figure S3.**
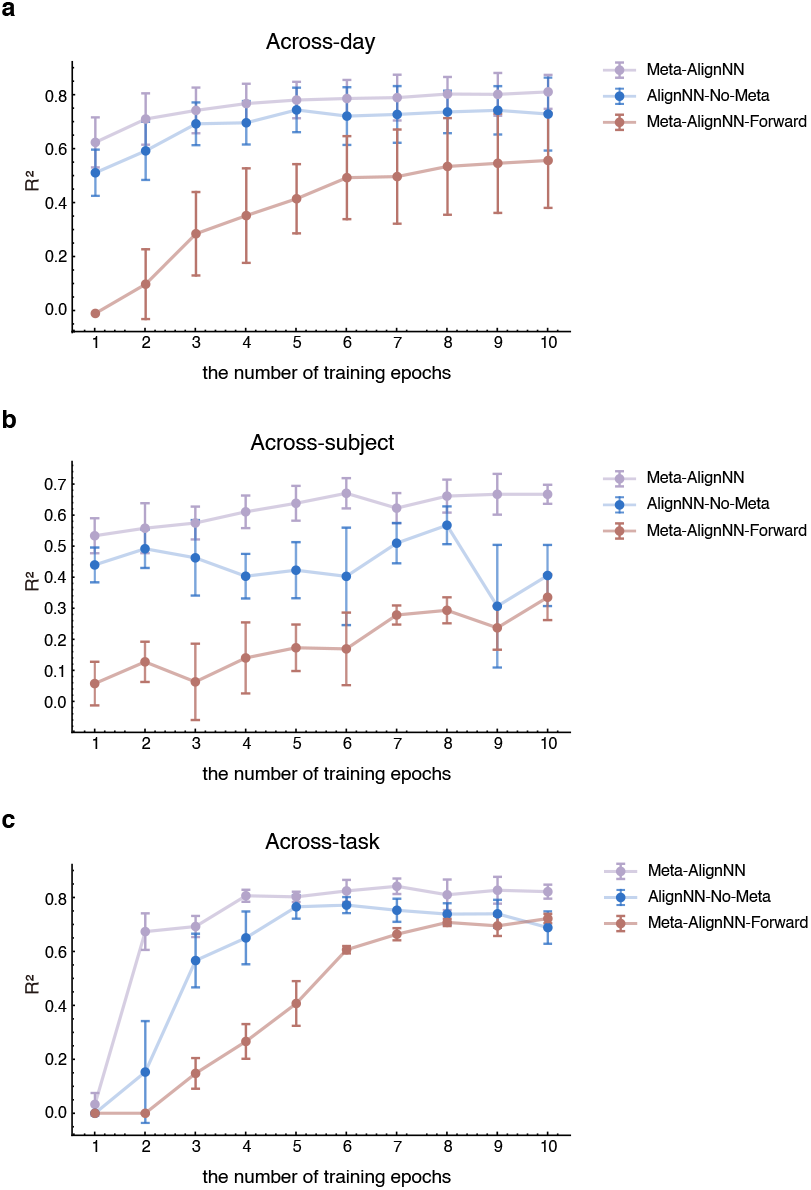
Ablation study. Behavioral decoding performance was quantified using mean R-squared values to contrast Meta-AlignNN, Meta-AlignNN-Forward, and AlignNN-No-Meta on held-out test sessions in three scenarios: across-day, across-subject, and across-task. X-axis represents the number of training epochs using a specific small amount of training data, after which the model is tested.

**Figure S4.**
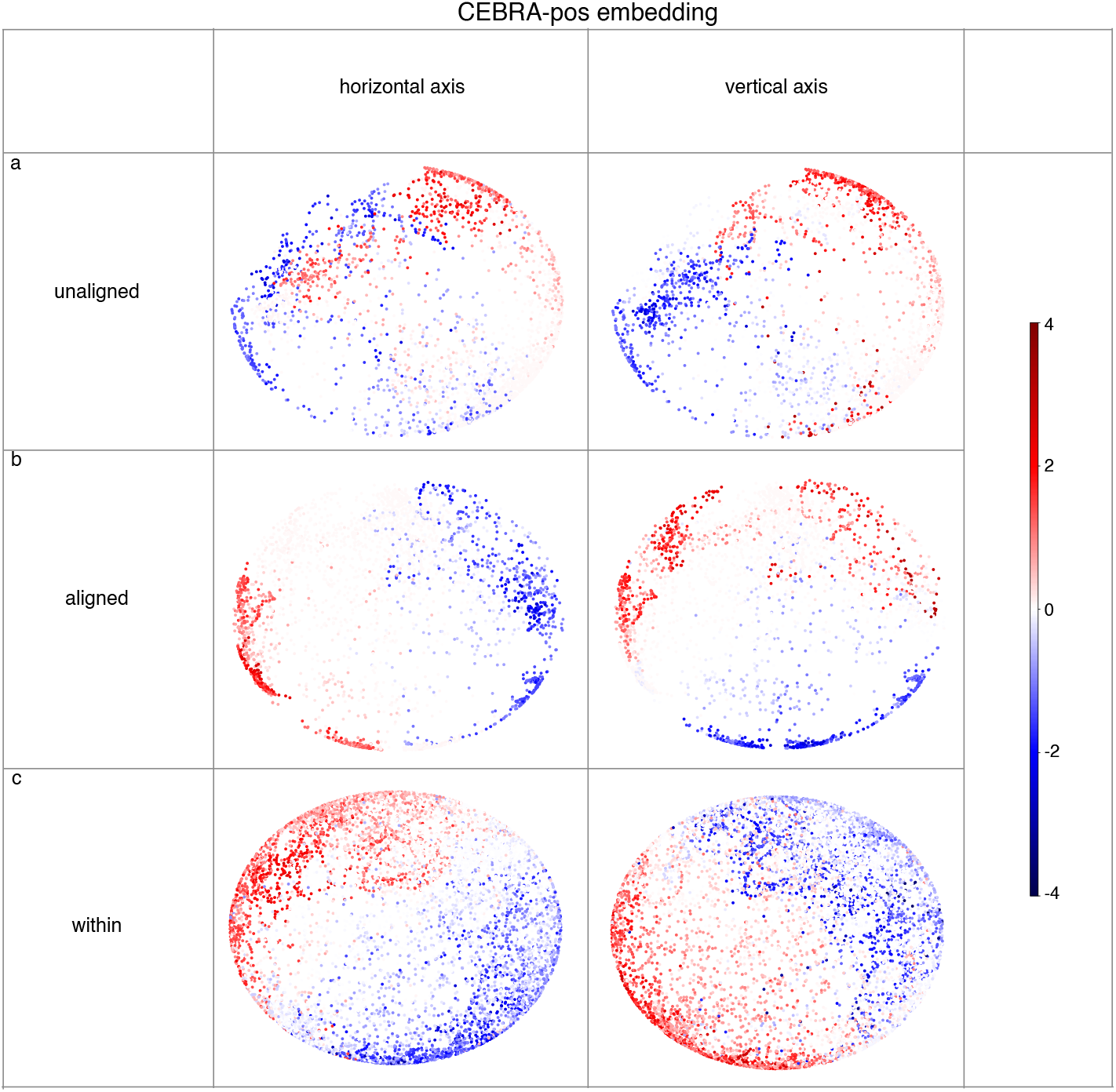
Comparison of CEBRA-Behavior embedding before and after alignment using Meta-AlignNN. By means of CEBRA-Behavior embedding with contrastive learning (Schneider et al., 2023), we want to compare the CEBRA-Behavior embedding of aligned neural activity with that of unaligned in js-8. Here, 3D CEBRA-Behavior embeddings represent neural-behavioral relationships, with points color-coded by normalized cursor position along the horizontal and vertical axes. In this figure, CEBRA-Behavior embedding was trained with position (x, y) of the cursor, and in each subplot, the left part represents color coded to the x position while the right part represents color coded to y position. The color bar denotes the Z-Score normalized value of cursor position relative to center point of screen. Each 3D point represents an embedding of neural activity extracted through CEBRA (Schneider et al., 2023), with the color intensity reflecting the cursor position at the corresponding moment. (**c**) shows the CEBRA-Behavior embedding of neural activity from one training session for the decoder. For the same test session, (**a**) shows the CEBRA-Behavior embedding of unaligned neural activity while (**b**) shows the CEBRA-Behavior embedding of aligned neural activity. It is not hard to understand that different CEBRA-Behavior embedding distribution indicates different difficulty for decoder to predict behavior movement. A “good” CEBRA-Behavior embedding distribution should look just like the color bar. The darkest red point should locate in one side of embedding space while darkest blue point should locate in the other side, and the lighter the color, the closer it is to the center. By comparing (**a**), (**b**), and (**c**), two conclusions can be readily drawn. 1. The distribution in (**c**) is better than (**a**), which indicates that with the shift and decay of electrodes, neural activity in unseen test sessions recorded by the same electrodes that was selected for past sessions is not kind for behavior decoding. 2. The distribution in (**b**) is better than (**a**), which indicates that the meta aligner successfully transform distribution of latter unseen neural activity into good neural pattern that is easy for behavior decoding. **a**, The CEBRA-behavior embedding is applied to unaligned neural activity from sessions that are months apart from sessions in which the electrodes were originally identified and selected. The signals recorded by the corresponding specific electrodes exhibit shifts and degradation. **b**, Meta-AlignNN stabilization: CEBRA-behavior embedding of aligned neural signals from matched recording sessions maintain good behavioral encoding topology. **c**, As a comparison, CEBRA-Behavior embedding was extracted from one of the sessions used to identify and select electrodes, and it reveals the intrinsic neural-behavior correspondence. Color bar: Normalized cursor coordinate along a single direction.

**Figure S5.**
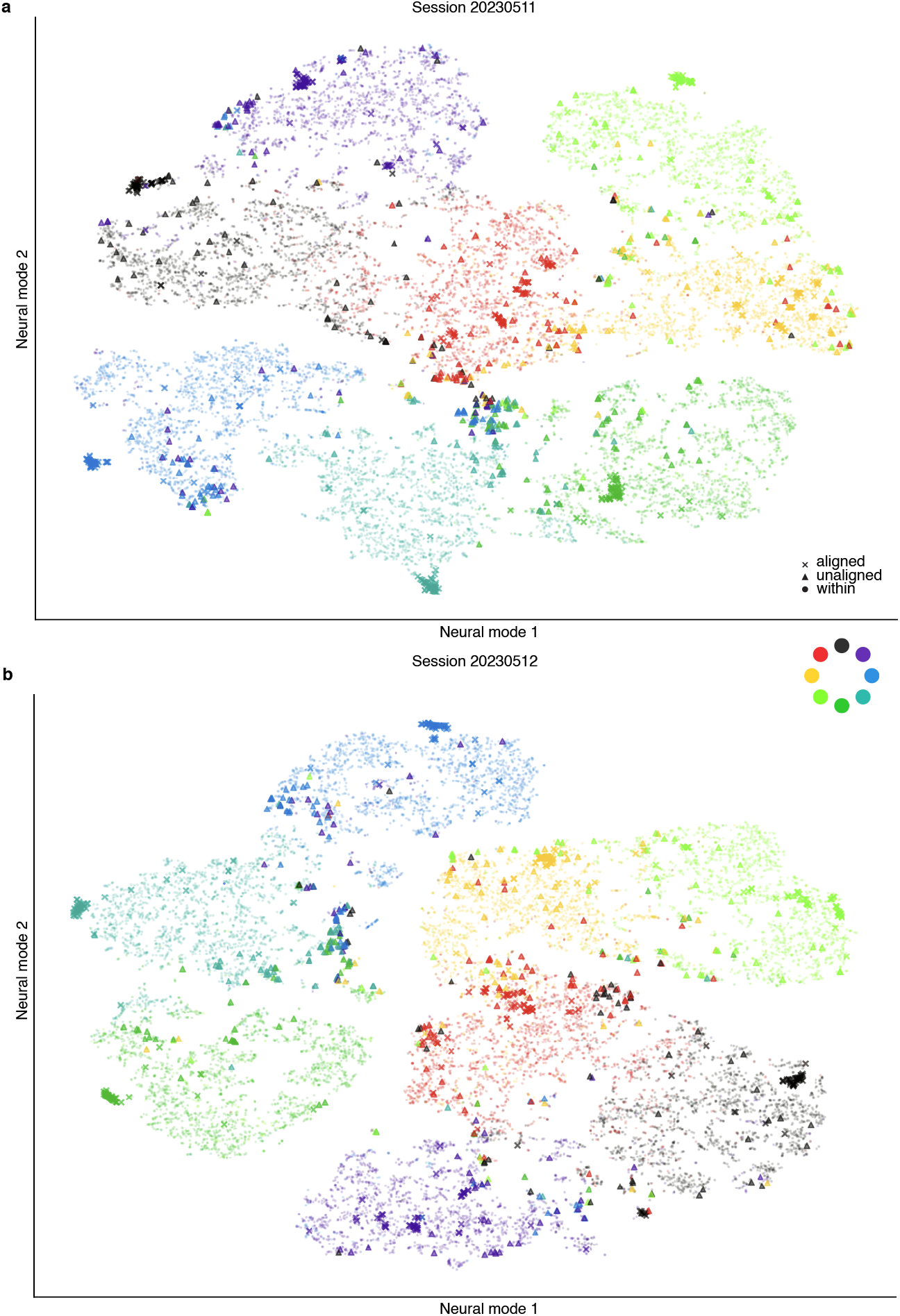
Comparison of neural latent space before and after alignment using Meta-AlignNN. This figure visualizes the t-SNE embeddings of Meta-AlignNN’s latent representations (the latent variable of the fixed decoder; see Methods for details) in two js-8 test sessions. Each color denotes specific target directions in the behavioral task and each dot (solid circle/triangle/x-shape) represents the embedding of latent representations for one trial. A different color mapping was used to represent the eight directions in order to make this figure clearer. Embeddings of the training trials for the fixed decoder are shown as tiny, semi-transparent solid circles, the unaligned test trials are shown as triangles, and the aligned test trials are shown as x-shape. As shown in this figure, application of Meta-AlignNN to any test session generates latent spaces exhibiting significant discriminability aligned with behavioral distinctions. In contrast, unaligned neural activities are mapped to incorrect latent space that is inconsistent with the latent space extracted by the fixed decoder, as you can see that large proportion of triangles are distributed in the region of circles with different color. This phenomenon intuitively explained what Meta-AlignNN does and why it works. In detail, eight separated clusters formed by circles corresponding to eight behavioral target direction denotes that the fixed decoder extracted an meaningful latent space from previous concatenating neural recordings over several days, while x-shape dots are almost distributed in corresponding color clusters means that the meta aligner successfully transform distribution of latter unseen neural activity into that of the selected former consecutive sessions and consequently result in the consistent latent representation by decoder. If the neural activity alignment fails, its effect will appear as shown by the triangles. We can draw the observation from this figure that incorrect distribution of unaligned latent embedding of test trials almost results from the adjacent target direction, e.g., red and light green triangles hold over 80 percent of error triangles distributed in yellow cluster. This demonstrates that behavioral similarity within a task correlates with neural representational similarity, posing inherent challenges for decoders to resolve such patterns under neural recording shift and change. However, few error distribution can be found with x-shape dots, which indicates that it is easy for the decoder to distinguish similar behaviors with different similar neural patterns, once the neural activity is aligned by meta aligner.

**Figure S6.**
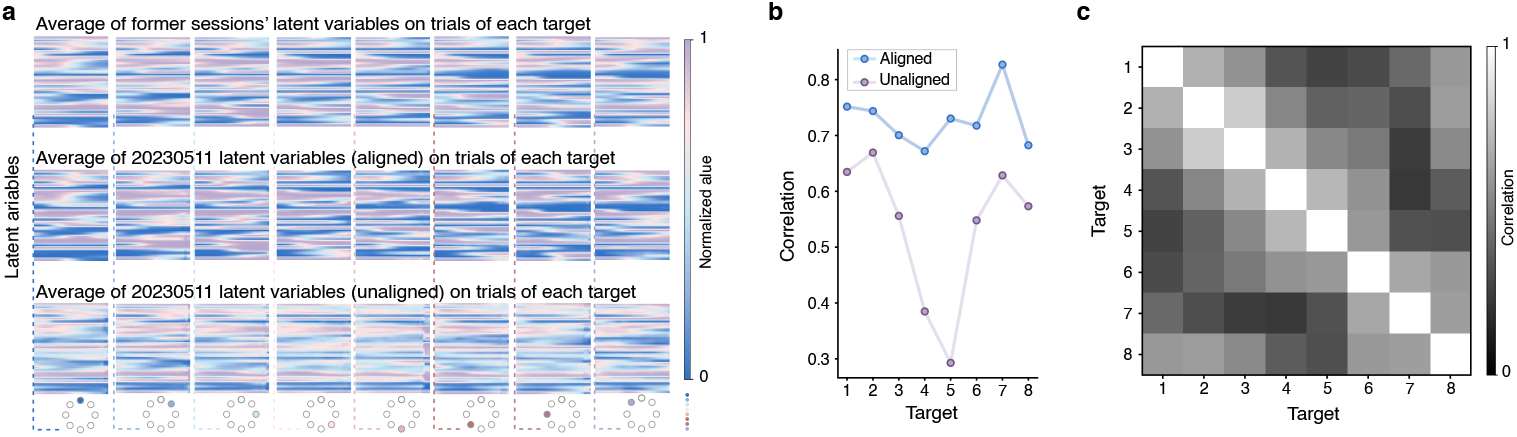
Neural latent variables. **a**, Mean 32D latent variables (1st row: Latent variables from former sessions; 2nd row: Aligned latent variables from session 20230511; 3rd row: Unaligned latent variables from from session 20230511) for corresponding target direction of js-8. Columns display trial-averaged latent variables for each of the eight target directions in js-8 (visualized below). **b**, Pairwise correlation analysis of latent variables across the eight target directions. **c**, Correlation matrix, computing the correlations between latent variables of different targets after alignment, e.g., matrix element (p, q) quantifies the Pearson correlation between latent representations of target p and target q post-alignment.

**Figure S7.**
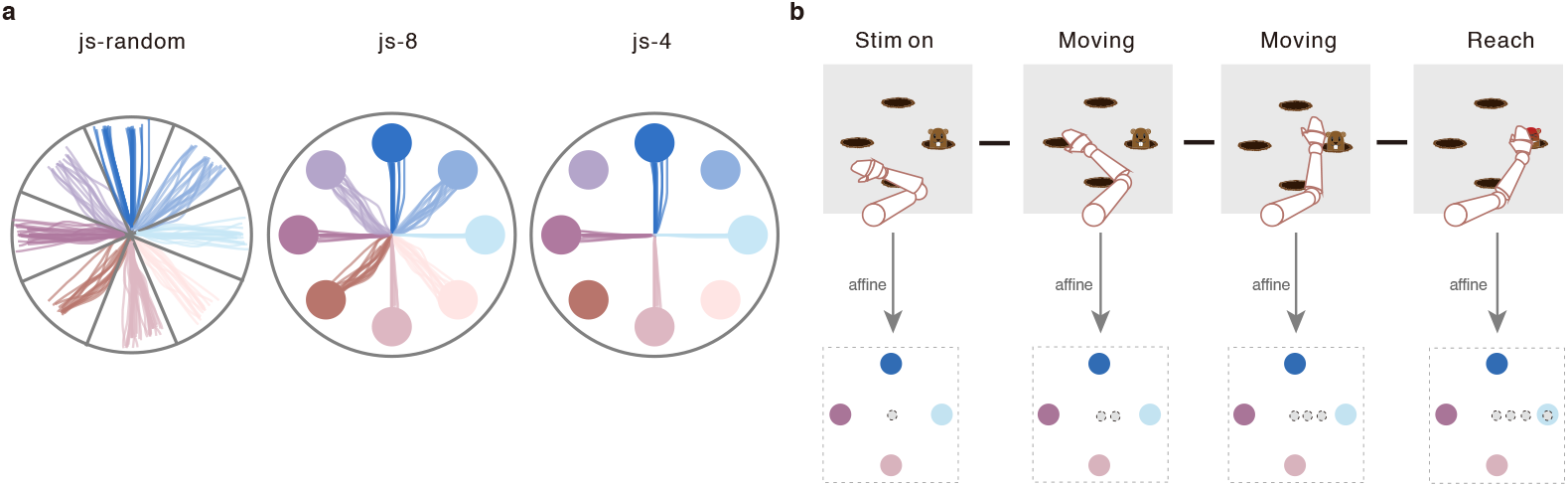
Variations across experimental tasks. **a**, Schematic movement trajectories of three tasks (js-random, js-8, js-4). **b**, Schematic diagram of affine transformation from classification to regression in wam-4.

**Figure S8.**
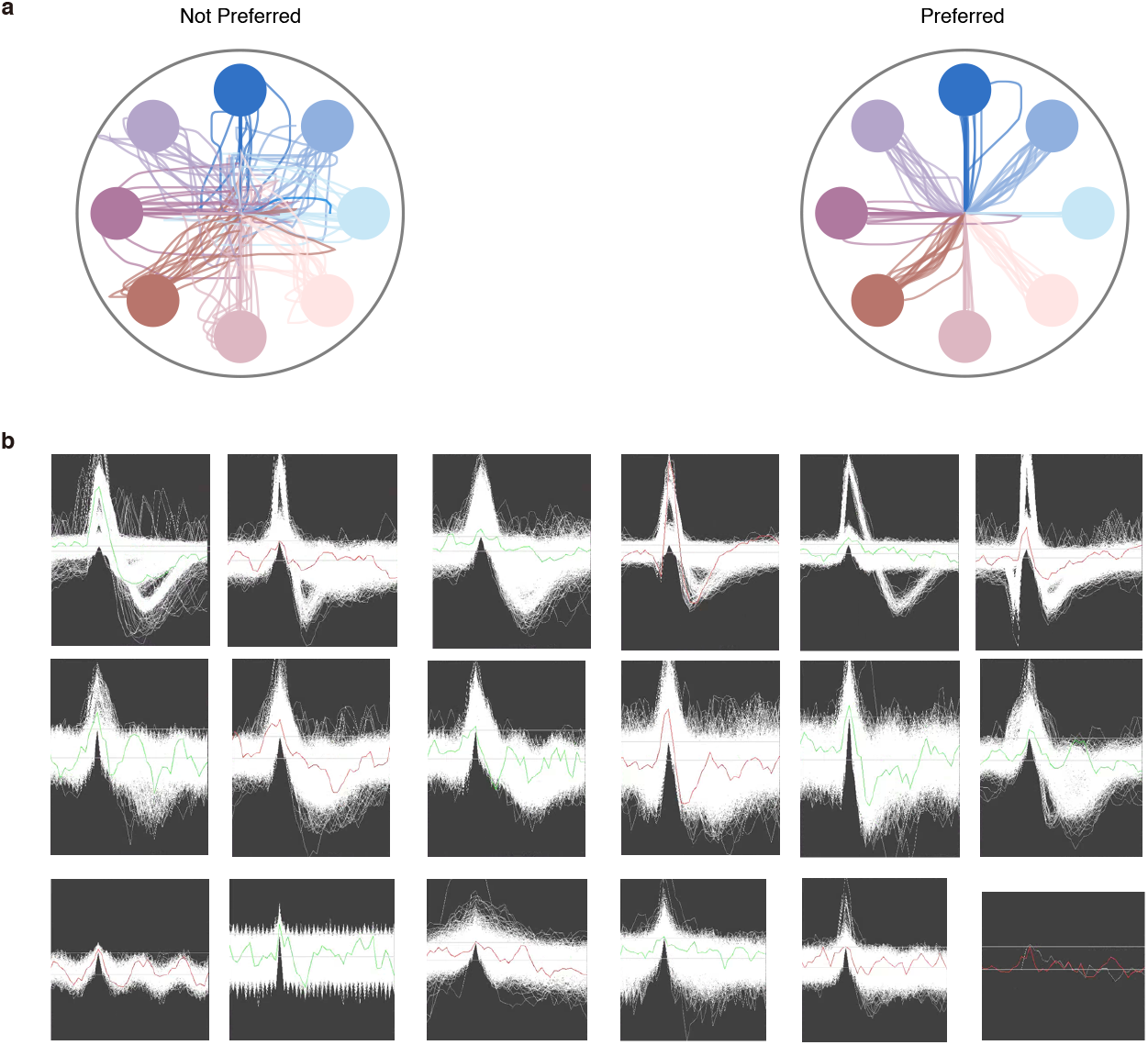
Identifying sessions and electrodes. **a** shows trial trajectories from two example sessions, and the session in which the monkey performed well (right) was selected. **b**, Multiunit threshold crossings from several example recording channels, with each panel representing one channel. Each panel contains a large number of spikes and green or red line denotes one random spike. Two horizontal lines indicate the detection threshold. We selected the channels in the first two rows and discarded those in the last row. Notably, the last panel in third row holds few recordings as corresponding electrode corrupted.

**Figure S9.**
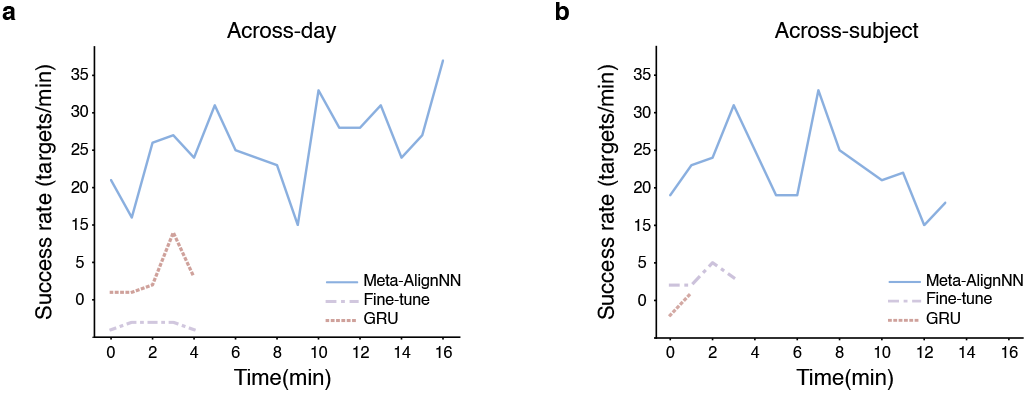
Online brain-control performance over longer timescales. **a**, In an example test session under the across-day scenario, online brain-control performance was compared among Meta-AlignNN, fine-tune, and GRU, measured by the number of targets hit per minute in js-8. Similarly, **b** shows online brain-control performance under the across-subject scenario (MK_RS <-> MK_NW).

**Figure S10.**
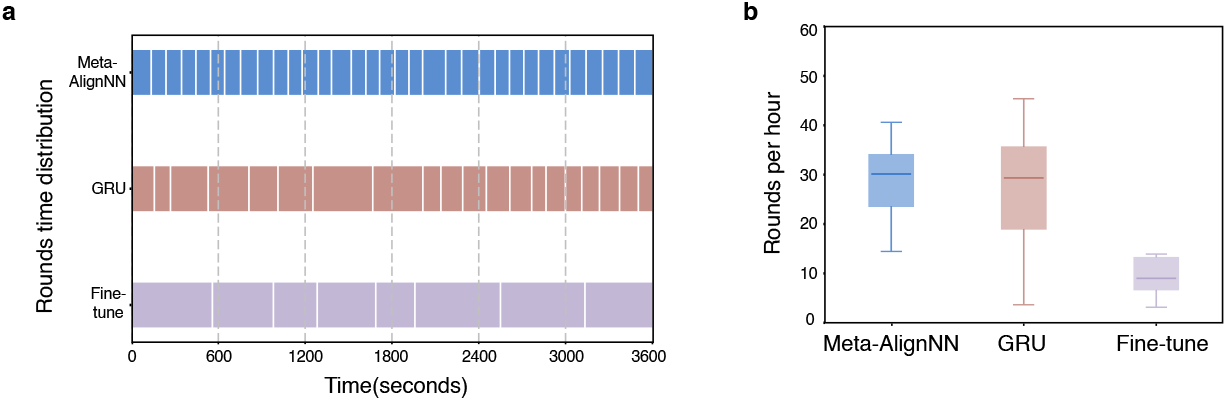
Online brain-control performance comparison in *Black Myth: Wukong*. In *Black Myth: Wukong*, completing a specific level (defeating two minions and one boss) is referred to as one round. **a**, Example comparison of the distribution of brain-control round completion times within one hour across the three models. **b**, Box plot statistics of online brain-control performance across multiple sessions.

## Reference

Bach, F. R. and Jordan, M. I. (2002). Kernel independent component analysis. Journal of machine learning research, 3(Jul):1–48.

Barbieri, R., Wilson, M. A., Frank, L. M., and Brown, E. N. (2005). An analysis of hippocampal spatio-temporal representations using a bayesian algorithm for neural spike train decoding. IEEE transactions on neural systems and rehabilitation engineering, 13(2):131–136.

Bishop, W., Chestek, C. C., Gilja, V., Nuyujukian, P., Foster, J. D., Ryu, S. I., Shenoy, K. V., and Yu, B. M. (2014). Self-recalibrating classifiers for intracortical brain-computer interfaces. Journal of Neural Engineering, 11(2):026001. doi: 10.1088/1741-2560/11/2/026001.

Buzsáki, G., Anastassiou, C. A., and Koch, C. (2012). The origin of extracellular fields and currents—eeg, ecog, lfp and spikes. Nature reviews neuroscience, 13(6):407–420.

Carmena, J. M., Lebedev, M. A., Crist, R. E., O’Doherty, J. E., Santucci, D. M., Dimitrov, D. F., Patil, P. G., Henriquez, C. S., and Nicolelis, M. A. L. (2003). Learning to control a brain–machine interface for reaching and grasping by primates. PLoS biology, 1(2):e42.

Chang, C.-C. and Lin, C.-J. (2011). Libsvm: a library for support vector machines. ACM transactions on intelligent systems and technology (TIST), 2(3):1–27.

Chestek, C. A., Batista, A. P., Santhanam, G., Byron, M. Y., Afshar, A., Cunningham, J. P., Gilja, V., Ryu, S. I., Churchland, M. M., and Shenoy, K. V. (2007). Single-neuron stability during repeated reaching in macaque premotor cortex. Journal of Neuroscience, 27(40): 10742–10750.

Chestek, C. A., Gilja, V., Nuyujukian, P., Foster, J. D., Fan, J. M., Kaufman, M. T., Churchland, M. M., Rivera-Alvidrez, Z., Cunningham, J. P., Ryu, S. I., and Shenoy, K. V. (2011). Longterm stability of neural prosthetic control signals from silicon cortical arrays in rhesus macaque motor cortex. Journal of Neural Engineering, 8(4):045005. doi: 10.1088/1741-2560/8/4/045005.

Cho, K., Van Merriënboer, B., Gulcehre, C., Bahdanau, D., Bougares, F., Schwenk, H., and Bengio, Y. (2014). Learning phrase representations using rnn encoder-decoder for statistical machine translation. arXiv preprint 1406.1078.

Churchland, M. M. and Shenoy, K. V. (2007). Temporal complexity and heterogeneity of single-neuron activity in premotor and motor cortex. Journal of Neurophysiology, 97(6):4235–4257. doi: 10.1152/jn.00095.2007.

Churchland, M. M., Cunningham, J. P., Kaufman, M. T., Foster, J. D., Nuyujukian, P., Ryu, S. I., and Shenoy, K. V. (2012). Neural population dynamics during reaching. Nature, 487(7405):51–56.

Dangi, S., Orsborn, A. L., Moorman, H. G., and Carmena, J. M. (2013). Design and analysis of closed-loop decoder adaptation algorithms for brain-machine interfaces. Neural Computation, 25(7):1693–1731. doi: 10.1162/neco_a_00460.

Degenhart, A. D., Bishop, W. E., Oby, E. R., Tyler-Kabara, E. C., Chase, S. M., Batista, A. P., and Yu, B. M. (2020). Stabilization of a brain-computer interface via the alignment of lowdimensional spaces of neural activity. Nature Biomedical Engineering, page 672–685. doi: 10.1038/s41551-020-0542-9.

Devlin, J., Chang, M.-W., Lee, K., and Toutanova, K. (2018). Bert: Pre-training of deep bidirectional transformers for language understanding. arXiv preprint 1810.04805.

Dickey, A. S., Suminski, A., Amit, Y., and Hatsopoulos, N. G. (2009). Single-unit stability using chronically implanted multielectrode arrays. Journal of neurophysiology, 102(2):1331–1339.

Downey, J. E., Schwed, N., Chase, S. M., Schwartz, A. B., and Collinger, J. L. (2018). Intracortical recording stability in human brain-computer interface users. Journal of Neural Engineering, 15(4):046016. doi: 10.1088/1741-2552/aab7a0.

Dyer, E. L., Gheshlaghi Azar, M., Perich, M. G., Fernandes, H. L., Naufel, S., Miller, L. E., and Körding, K. P. (2017). A cryptography-based approach for movement decoding. Nature biomedical engineering, 1(12):967–976.

Farshchian, A., Gallego, J., Cohen, J., Bengio, Y., Miller, L., and Solla, S. (2018). Adversarial domain adaptation for stable brain-machine interfaces. International Conference on Learning Representations,International Conference on Learning Representations.

Finn, C., Abbeel, P., and Levine, S. Model-agnostic meta-learning for fast adaptation of deep networks. In International conference on machine learning, pages 1126–1135. PMLR, (2017).

Flint, R. D., Scheid, M. R., Wright, Z. A., Solla, S. A., and Slutzky, M. W. (2016). Long-term stability of motor cortical activity: implications for brain machine interfaces and optimal feedback control. Journal of Neuroscience, 36(12):3623–3632.

Fraser, G. W. and Schwartz, A. B. (2012). Recording from the same neurons chronically in motor cortex. Journal of neurophysiology, 107(7):1970–1978.

Gallego, J. A., Perich, M. G., Naufel, S. N., Ethier, C., Solla, S. A., and Miller, L. E. (2018). Cortical population activity within a preserved neural manifold underlies multiple motor behaviors. Nature communications, 9(1):4233.

Gallego, J. A., Perich, M. G., Chowdhury, R. H., Solla, S. A., and Miller, L. E. (2020). Longterm stability of cortical population dynamics underlying consistent behavior. Nature neuroscience, 23(2):260–270.

Ganin, Y., Ustinova, E., Ajakan, H., Germain, P., Larochelle, H., Laviolette, F., Marchand, M., and Lempitsky, V. (2016). Domain-adversarial training of neural networks. The journal of machine learning research, 17(1):2096–2030.

Gilja, V., Nuyujukian, P., Chestek, C. A., Cunningham, J. P., Yu, B. M., Fan, J. M., Churchland, M. M., Kaufman, M. T., Kao, J. C., Ryu, S. I., et al. (2012). A high-performance neural prosthesis enabled by control algorithm design. Nature neuroscience, 15(12):1752–1757.

Goodfellow, I., Bengio, Y., and Courville, A. Deep learning. MIT press, (2016).

Goodfellow, I. J., Pouget-Abadie, J., Mirza, M., Xu, B., Warde-Farley, D., Ozair, S., Courville, A., and Bengio, Y. (2014). Generative adversarial nets. Advances in neural information processing systems, 27.

Jude, J., Perich, M. G., Miller, L. E., and Hennig, M. H. (2022). Robust alignment of cross-session recordings of neural population activity by behaviour via unsupervised domain adaptation. arXiv preprint 2202.06159.

Kao, J. C., Ryu, S. I., and Shenoy, K. V. (2017). Leveraging neural dynamics to extend functional lifetime of brain-machine interfaces. Scientific reports, 7(1):7395.

Kingma, D. P. and Ba, J. (2014). Adam: A method for stochastic optimization. arXiv preprint 1412.6980.

Kloosterman, F., Layton, S. P., Chen, Z., and Wilson, M. A. (2014). Bayesian decoding using unsorted spikes in the rat hippocampus. Journal of neurophysiology, 111(1):217–227.

Natekin, A. and Knoll, A. (2013). Gradient boosting machines, a tutorial. Frontiers in neurorobotics, 7:21.

Nichol, A., Achiam, J., and Schulman, J. (2018). On first-order meta-learning algorithms. arXiv preprint 1803.02999.

Nuyujukian, P., Kao, J. C., Fan, J. M., Stavisky, S. D., Ryu, S. I., and Shenoy, K. V. (2014). Performance sustaining intracortical neural prostheses. Journal of neural engineering, 11(6):066003.

O’Doherty, J. E., Cardoso, M. M. B., Makin, J. G., and Sabes, P. N. Nonhuman primate reaching with multichannel sensorimotor cortex electrophysiology, (2017). URL 10.5281/zenodo.788569.

Orbach, J. (1962). Principles of neurodynamics. perceptrons and the theory of brain mechanisms. Archives of General Psychiatry, 7(3):218–219.

Orsborn, A. L., Dangi, S., Moorman, H. G., and Carmena, J. M. (2012). Closed-loop decoder adaptation on intermediate time-scales facilitates rapid bmi performance improvements independent of decoder initialization conditions. IEEE Transactions on Neural Systems and Rehabilitation Engineering, 20(4):468–477. doi: 10.1109/tnsre.2012.2185066.

Pandarinath, C., O’Shea, D. J., Collins, J., Jozefowicz, R., Stavisky, S. D., Kao, J. C., Trautmann, E. M., Kaufman, M. T., Ryu, S. I., Hochberg, L. R., et al. (2018). Inferring singletrial neural population dynamics using sequential auto-encoders. Nature methods, 15 (10):805–815.

Perge, J. A., Homer, M. L., Malik, W. Q., Cash, S., Eskandar, E., Friehs, G., Donoghue, J. P., and Hochberg, L. R. (2013). Intra-day signal instabilities affect decoding performance in an intracortical neural interface system. Journal of Neural Engineering, 10(3):036004. doi: 10.1088/1741-2560/10/3/036004.

Pohlmeyer, E. A., Solla, S. A., Perreault, E. J., and Miller, L. E. (2007). Prediction of upper limb muscle activity from motor cortical discharge during reaching. Journal of neural engineering, 4(4):369.

Rumelhart, D. E., Hinton, G. E., and Williams, R. J. (1986). Learning representations by back-propagating errors. nature, 323(6088):533–536.

Sadtler, P. T., Quick, K. M., Golub, M. D., Chase, S. M., Ryu, S. I., Tyler-Kabara, E. C., Yu, B. M., and Batista, A. P. (2014). Neural constraints on learning. Nature, 512(7515):423–426.

Safaie, M., Chang, J. C., Park, J., Miller, L. E., Dudman, J. T., Perich, M. G., and Gallego, J. A. (2023). Preserved neural dynamics across animals performing similar behaviour. Nature, 623(7988):765–771.

Santhanam, G., Linderman, M. D., Gilja, V., Afshar, A., Ryu, S. I., Meng, T. H., and Shenoy, K. V. (2007). Hermesb: A continuous neural recording system for freely behaving primates. IEEE Transactions on Biomedical Engineering, page 2037–2050. doi: 10.1109/tbme.2007.895753.

Schaul, T. and Schmidhuber, J. (2010). Metalearning. Scholarpedia, 5(6):4650. doi: 10.4249/scholarpedia.4650. revision #91489.

Schmidhuber, J. Evolutionary principles in self-referential learning, or on learning how to learn: the meta-meta-… hook. PhD thesis, Technische Universität München, (1987).

Shenoy, K. V., Sahani, M., and Churchland, M. M. (2013). Cortical control of arm movements: a dynamical systems perspective. Annual review of neuroscience, 36(1):337–359.

Smola, A. J. and Schölkopf, B. (2004). A tutorial on support vector regression. Statistics and computing, 14:199–222.

Stevenson, I. H., Cherian, A., London, B. M., Sachs, N. A., Lindberg, E., Reimer, J., Slutzky, M. W., Hatsopoulos, N. G., Miller, L. E., and Kording, K. P. (2011). Statistical assessment of the stability of neural movement representations. Journal of neurophysiology, 106(2):764–774.

Sussillo, D., Stavisky, S. D., Kao, J. C., Ryu, S. I., and Shenoy, K. V. (2016). Making brain– machine interfaces robust to future neural variability. Nature communications, 7(1):13749.

Tian, Y., Zhao, X., and Huang, W. (2022). Meta-learning approaches for learning-to-learn in deep learning: A survey. Neurocomputing, 494:203–223.

Tolias, A. S., Ecker, A. S., Siapas, A. G., Hoenselaar, A., Keliris, G. A., and Logothetis, N. K. (2007). Recording chronically from the same neurons in awake, behaving primates. Journal of neurophysiology, 98(6):3780–3790.

Vaswani, A., Shazeer, N., Parmar, N., Uszkoreit, J., Jones, L., Gomez, A. N., Kaiser, Ł., and Polosukhin, I. (2017). Attention is all you need. Advances in neural information processing systems, 30.

Wen, S., Yin, A., Furlanello, T., Perich, M. G., Miller, L. E., and Itti, L. (2023). Rapid adaptation of brain–computer interfaces to new neuronal ensembles or participants via generative modelling. Nature biomedical engineering, 7(4):546–558.

Willett, F. R., Avansino, D. T., Hochberg, L. R., Henderson, J. M., and Shenoy, K. V. Highperformance brain-to-text communication via imagined handwriting. (2020). URL 10.1101/2020.07.01.183384.

Wu, W. and Hatsopoulos, N. G. (2008). Real-time decoding of nonstationary neural activity in motor cortex. IEEE Transactions on Neural Systems and Rehabilitation Engineering, 16(3):213–222. doi: 10.1109/tnsre.2008.922679.

Wu, W., Gao, Y., Bienenstock, E., Donoghue, J. P., and Black, M. J. (2006). Bayesian population decoding of motor cortical activity using a kalman filter. Neural computation, 18(1):80–118.

Yu, B. M., Cunningham, J. P., Santhanam, G., Ryu, S., Shenoy, K. V., and Sahani, M. (2008). Gaussian-process factor analysis for low-dimensional single-trial analysis of neural population activity. Advances in neural information processing systems, 21.

Zhang, Y. and Chase, S. M. A stabilized dual kalman filter for adaptive tracking of braincomputer interface decoding parameters. In 2013 35th Annual International Conference of the IEEE Engineering in Medicine and Biology Society (EMBC), pages 7100–7103. IEEE, (2013).

## Supplementary Reference

Schneider, S., Lee, J. H., and Mathis, M. W. (2023). Learnable latent embeddings for joint behavioural and neural analysis. Nature, pages 1–9.

